# Toward industrial C8 production: Oxygen intrusion drives renewable *n*-caprylate production from ethanol and acetate *via* intermediate metabolite production

**DOI:** 10.1101/2024.07.12.603245

**Authors:** Kurt Gemeinhardt, Byoung Seung Jeon, Jean Nepomuscene Ntihuga, Han Wang, Caroline Schlaiß, Timo N. Lucas, Irina Bessarab, Nicolas Nalpas, Nanqing Zhou, Joseph G. Usack, Daniel H. Huson, Rohan B. H. Williams, Boris Maček, Ludmilla Aristilde, Largus T. Angenent

## Abstract

Previous bioreactor studies achieved high volumetric *n*-caprylate (*i.e., n*-octanoate) production rates and selectivities from ethanol and acetate with chain-elongating microbiomes. However, the metabolic pathways from the substrates to *n*-caprylate synthesis were unclear. We operated two *n*-caprylate-producing upflow bioreactors with a synthetic medium to study the underlying metabolic pathways. The operating period exceeded 2.5 years, with a peak volumetric *n*-caprylate production rate of 190 ± 8.4 mmol C L^-1^ d^-1^ (0.14 g L^-1^ h^-1^). We identified oxygen availability as a critical performance parameter, facilitating intermediate metabolite production from ethanol. Bottle experiments in the presence and absence of oxygen with ^13^C-labeled ethanol suggest acetyl-coenzyme A-based derived production of *n*-butyrate (*i.e., n*-butanoate), *n*-caproate (*i.e., n*-hexanoate), and *n*-caprylate. Here, we postulate a trophic hierarchy within the bioreactor microbiomes based on metagenomics, metaproteomics, and metabolomics data, as well as experiments with a *Clostridium kluyveri* isolate. First, the aerobic bacterium *Pseudoclavibacter caeni* and the facultative anaerobic fungus *Cyberlindnera jadinii* converted part of the ethanol pool into the intermediate metabolites succinate, lactate, and pyroglutamate. Second, the strict anaerobic *C. kluyveri* elongated acetate with the residual ethanol to *n*-butyrate. Third, *Caproicibacter fermentans* and *Oscillibacter valericigenes* elongated *n*-butyrate with the intermediate metabolites to *n*-caproate and then to *n*-caprylate. Among the carbon chain-elongating pathways of carboxylates, the tricarboxylic acid cycle and the reverse ß-oxidation pathways showed a positive correlation with *n*-caprylate production. The results of this study inspire the realization of a chain-elongating production platform with separately controlled aerobic and anaerobic stages to produce *n*-caprylate renewably as an attractive chemical from ethanol and acetate as substrates.

**Broader context:** Next to renewable electric energy, carbon-based chemicals have to be produced sustainably and independently from fossil sources. To meet this goal, we must expand the portfolio of bio-based conversion technologies on an industrial scale to cover as many target chemicals as possible. We explore the bioprocess of chain elongation to provide medium-chain carboxylates that can function as future platform chemicals in the circular economy. The most valuable medium-chain carboxylate produced with chain elongation is *n*-caprylate (*i.e., n*-octanoate). This molecule with eight carbon atoms in a row (C8) is challenging to produce renewably for the chemical industry. Previous reports elucidated that elevated ethanol-to-acetate ratios, which are found in syngas-fermentation effluent, stimulated *n*-caprylate production. Until now, studies have suggested that chain elongation from high concentrations of ethanol and acetate is a fully anaerobic process. We refine this view by showing a trophic hierarchy of aerobic and anaerobic microbes capable of facilitating this process. Appropriate oxygen supplementation enables the synthesis of succinate, lactate, and pyroglutamate that permit high-rate chain elongation to *n*-caprylate under anaerobic conditions. Given these results, future research should focus on the segregated study of aerobic and anaerobic microbes to further enhance the process performance to produce *n*-caprylate renewably at an industrial scale.

## Introduction

Microbial chain elongation is a bioprocess for medium-chain carboxylate production that expands the portfolio of sustainably sourced platform chemicals for the circular economy.^1-3^ Throughout the text, we refer to carboxylates as their combined undissociated and dissociated species with one carboxyl group. Medium-chain carboxylates (*i.e*., carbon skeleton of C6 to C12) are of industrial interest with applications such as antimicrobials, fragrance precursors, and drug delivery supplements.^4, 5^ Most importantly, they can be building blocks for synthesizing additional chemicals.^6^ The possible chemical derivatives are plentiful and they include ketones, alcohols, fatty alcohols, fatty acid alkyl esters, and longer-chain alkanes *via* Kolbe electrolysis.^7, 8^ In the case of *n*-caprylate, Kolbe electrolysis yields *n*-tetradecane, which is a drop-in fuel for aviation.^9, 10^

Chain elongators are anaerobic bacteria that produce medium-chain carboxylates such as *n*-caproate (six carbon atoms; C6; *n*-hexanoate) and *n*-caprylate (C8; *n*-octanoate) from short-chain carboxylates, such as acetate (C2) and *n*-butyrate (C4; *n*-butanoate), often with ethanol (C2) as an electron donor.^1, 11, 12^ Chain elongation with open cultures produces an array of short to medium-chain carboxylates, with C8 as the longest carbon chain, at reasonable rates.^13^ *n*-Caprylate has a higher monetary value than shorter-chain carboxylates. Moreover, as the length of the carbon chain increases, the effect of the hydrophobic tail predominates over the hydrophilic carboxyl group, facilitating extraction from water. *n*-Caprylate production from palm and coconut oil on an industrial scale is unsustainable. ^14^ The extraction by fractionation from a small portion of these oils is scarce, hindering global market development.^15^ Because of the superior properties of *n*-caprylate, we explore strategies to boost the production rate and selectivity (% specific product per total substrate fed) of chain-elongating bioprocesses. The chemical industry is looking for an economically promising method to produce C8 compounds renewably.^16^

To produce *n*-caprylate, the activity of methanogenic archaea must be controlled; otherwise, the reducing power of anaerobic microbiomes channels electrons into the most reduced carbon species – methane.^17^ The two relevant types of methanogens are acetoclastic and hydrogenotrophic archaea. Acetoclastic methanogens convert acetate into carbon dioxide and methane, depriving the chain elongators of their substrate. A mildly acidic pH of 5.0-5.5 strongly inhibits acetoclastic methanogens.^13^ Hydrogenotrophic methanogens convert molecular hydrogen and carbon dioxide into methane. On the one hand, reducing the hydrogen partial pressure below 0.02 atm leads to excessive ethanol oxidation, which can reduce the production rates of medium-chain carboxylate in anaerobic systems.^18^ On the other hand, hydrogen is a side product of chain elongation. Therefore, lowering the hydrogen partial pressure can increase the volumetric *n*-caprylate production rate by side-product removal.^19^

Few open-culture studies observed high *n*-caprylate selectivities and almost all with ethanol and acetate as the substrate.^20-23^ The flux of successive metabolic reactions from ethanol and acetate to *n*-caprylate is speculative due to the challenging task of obtaining information from bioreactor microbiomes.^24^ Previous studies suggest the reverse ß-oxidation pathway as the primary pathway to produce *n*-caprylate.^19, 25, 26^ However, the most commonly used model microbe of the reverse ß-oxidation pathway, which is *Clostridium kluyveri*, can only produce minor amounts of *n*-caprylate through unspecific enzyme reactions.^27-29^

Here, we investigated the long-term *n*-caprylate production in two bioreactors fed with a defined medium, containing high concentrations of ethanol (∼13.5 g L^-1^) and acetate (∼3 g L^-1^) while omitting yeast extract. Previous studies have elucidated the process setup and established the ethanol-to-acetate ratio as an essential parameter to sustain high volumetric *n*-caprylate production rates.^21, 22^ Here, we introduce oxygen supplementation as a crucial regulator of *n*-caprylate production. We used bottle experiments with a chain-elongating microbiome and stable-isotope tracing to monitor the molecular assembly of *n*-butyrate, *n*-caproate, and *n*-caprylate. Furthermore, a metagenomics analysis revealed the key species involved in *n*-caprylate production. The isolation of a previously undescribed *C. kluyveri* strain elucidated its ecological function in the bioreactor food web. Comparative metaproteomics revealed the active metabolic pathways in the microbiome. With metabolomics analysis, we determined pertinent metabolites as intermediates toward *n*-caprylate production that we monitored in additional bottle experiments.

### Experimental

We summarized the experimental methods in the main text. The electronic supplementary information (ESI †) comprises a detailed description of the used methods, materials, and instrumentation, including the model and manufacturer of equipment.

### Initial bioprocess configuration

We set up two independently and continuously fed upflow bioreactors with fermentation-broth recirculation through a membrane-based liquid-liquid extraction (*i.e*., pertraction) system (Bioreactor setup and operation, ESI †). Each bioreactor vessel consisted of double-walled glass in a vertical cylindric shape (125 cm height). A thermostatically controlled system maintained the temperature in the interior at 30 °C, and automatic acid-addition pumps adjusted the pH of the fermentation broth to 5.5. We filled one bioreactor vessel with plastic, wheel-shaped packing material, designated as the anaerobic filter (AF) reactor. We operated the second bioreactor as an upflow anaerobic sludge blanket (UASB) reactor without packing material to promote biomass accumulation as granules at the bottom of the vessel (**Fig. S1A-B**, ESI †).

The extraction systems of both bioreactors for extracting medium chain carboxylates from the fermentation broth were identical.^13^ A peristaltic pump recycled the fermentation broth (pH = 5.5) between the bioreactor vessel and the extraction system, consisting of two hollow-fiber membrane contactors (Extraction system, ESI †). A filter module with a wet volume of 0.8 L was connected in-line between the bioreactor vessel and extraction system to prevent membrane fouling for the first 732 days. The first membrane contactor created a liquid-liquid interface between the fermentation broth and a hydrophobic solvent. Hydrophobic products (medium-chain carboxylates) diffused into the hydrophobic solvent. Another peristaltic pump recycled the hydrophobic solvent between the first and the second membrane contactors. In the second membrane contactor, a pH-induced concentration gradient between undissociated and dissociated carboxylates caused the directed diffusion of carboxylates into an alkaline extraction solution (pH = 9.0).

### Operating periods I-IV

The reported operating period was preceded by a pre-operating period of 230 days (data not shown). During the pre-operating period, we determined the correct process parameters to operate the bioprocess stably (*e.g*., the recycling flow rate of the fermentation broth, the hydrophobic solvent, the alkaline extraction solution, and the substrate concentration). The reported operating period was 1019 days (∼2.8 years). We designated the first reported day as Day 0. Throughout the operating period, we applied several configurational modifications to the bioreactors regarding oxygen availability. Based on the modifications, we divided the reported operating timeframe into four distinct periods: **(1)** a stable *n*-caprylate production-rate period (passive oxygen intrusion into the system); **(2)** a transition period (passive oxygen intrusion stepwise limited); **(3)** a minimal *n*-caprylate production-rate period (no oxygen intrusion into the system); and **(4)** a fluctuating *n*-caprylate production-rate period (active oxygen supply). During the end of Period I and throughout Period II and III, we measured dissolved oxygen concentrations in the fermentation broth at different locations in the recycle line with non-invasive oxygen sensor spots (Analytical procedures, ESI †). Changes in the construction of the bioreactors had consequences regarding the wet working volume, hydraulic retention time, and thus the total organic loading rate (**Table S1** and **Eq. S1-4**, ESI †).

During Period I (Day 0-539), we continuously supplied a defined bioreactor medium (*i.e*., medium) to the bioreactors from a 4 L-vessel that was open to the atmosphere. On Day 174, we started supplying reducing agents to the bioreactors. First, we supplemented L-cysteine at a concentration of 0.3 g L^-1^ directly to the medium. After noticing a sharp decline in performance, we switched to supplying a different reducing agent from a separate 0.5-L bottle. A peristaltic pump, which was connected to a time switch, pulsed the reducing agent solution directly into the fermentation broth. The dosing cycle was 1 min every 2.4 h, totaling 10 min d^-1^. Additionally, we changed the reducing agent to sodium sulfide at an application rate of 0.2 g L^-1^ d^-1^ on Day 194 and lowered the rate to 0.02 g L^-1^ d^-1^ on Day 209. The decreased sodium sulfide application rate was maintained for the rest of the operating period except for Day 419-437, when we reiterated the L-cysteine spike into the medium.

During Period II (Day 550-732), we performed successive constructional modifications to limit oxygen intrusion into the bioreactors. On Day 550, we connected a swan-neck airlock to the effluent line. The fermentation broth exited the bioreactor vessel *via* a natural hydraulic flow through the swan-neck. Oxygen penetration into the system was limited to the liquid surface area inside the swan-neck. Next, we started continuously sparging nitrogen gas into the headspace of the bioreactors on Day 614 and into the medium tank on Day 645. On Day 701, we exchanged all the slightly gas-transmissible plastic tubing for gas-impermeable metal tubing. The last constructional change on Day 733, marking the end of Period II, was the removal of the filter modules, which were situated in line between the bioreactor vessel and extraction system.

During Period III (Day 733-861), on Day 747, we reduced the substrate concentration to 80% of the original value and on Day 767 to 60%. Further adjustments during Periods III and IV (Day 862-1019) concerned the introduction of reactive gases through the inlet line at the bottom of the bioreactor columns. First, we sparged N_2_/H_2_ (95/5% v/v) at a flow rate of 0.3 ± 0.1 ml h^-1^ during Day 784-846. Second, we changed the sparged gas to air (∼21% O_2_ v/v) with a flow rate of 11.8 ± 0.90 mL h^-1^ during Day 862-1003. As we noticed an increase in process performance, we increased the substrate concentration back to 80% on Day 971 and the airflow rate to 17.1 ± 1.5 ml h^-1^ from Day 1004 until the operating end (Day 1019).

### Medium and inoculum

We fed medium to the bioreactors, which previous studies established for chain-elongating microbiomes (Defined bioreactor medium, ESI †).^21, 22, 30, 31^ Modifications involved omitting yeast extract and including 3 g L^-1^ sodium bicarbonate. As the substrate, we supplemented ethanol and acetate at concentrations of 13.42 g L^-1^ (∼600 mM C) and 3.12 g L^-1^ (∼100 mM C), resulting in a ratio of 6:1 (mol/mol, ethanol:acetate) (**Table S1**, ESI †). We prepared the medium aerobically without sterilization, adding hydrochloric acid or sodium hydroxide to adjust to pH 5. On the first day of the pre-operating period, we inoculated both bioreactors with a 2-mL glycerol stock (0.04% v/v) that had been stored at -20 °C. The glycerol stock contained a well-studied chain-elongating bioreactor microbiome.^22^ The previous bioreactor was a completely stirred tank reactor (CSTR) with product extraction and was fed ethanol and acetate-rich medium for a year. Because we did not sterilize the defined growth medium during the operating period, and the bioreactor system was open to the environment (open culture), external microbial species entered the system, possibly changing the microbial community.

### Bottle experiments with and without stable-isotope tracing

We conducted bottle experiments with stable-isotope tracing with aerobic and anaerobic starting conditions (*i.e*., aerobic and anaerobic treatment) in 50-mL serum bottles. The wet working volume was 10 mL, and the headspace was 40 mL. We modified the medium for the bottle experiments by adding 0.5 g L^-1^ L-cysteine hydrochloride, replacing sodium bicarbonate with 3 g L^-1^ sodium carbonate, and omitting 2-(*n*-morpholino)ethanesulfonic acid buffer. To measure differences in the isotopic composition of products, we changed the substrate to 4.5 g L^-1^ (200 mM C) [1,2-^13^C_2_] ethanol without acetate. We capped the bottles with butyl stoppers and sealed them with aluminum caps. We prepared the aerobic bottles under ambient air conditions and flushed the anaerobic bottles with nitrogen gas for 30 min, displacing the contained oxygen before inoculation. The working volume consisted of 9.5 mL modified medium and 0.5 mL inoculum (5% v/v). The inoculum consisted of fresh fermentation broth from the AF reactor, which we directly transferred to the bottles on Day 412 when *n*-caprylate production rates were high. The bottle experiments lasted 10 days, and except for sampling periods, we stored the bottles in a 30 °C incubator without shaking. Daily, we monitored the pH, cell density, ethanol, carboxylate concentrations, and the isotopic compositions of produced carboxylates (Analytical procedures, ESI †). We conducted the experiments for the aerobic and anaerobic conditions in duplicates, and the given values represent averages. To complement the stable-isotope tracing experiments, we conducted additional bottle experiments with slightly changed media compositions, using unlabeled substrate, and thawed biomass as inoculum, and including metabolite concentration measurements of succinate, lactate, pyroglutamate, and pyruvate (Bottle experiments without stable-isotope tracing, ESI †).

### Chemical analyses and calculations

We collected liquid samples (1 mL) from the fermentation broth and alkaline extraction solution of both bioreactors daily or every other day. The fermentation broth samples originated from a sampling port at one-fifth (25 cm) of the height of the bioreactor column. The alkaline extraction solution samples were from a T-shaped sampling adaptor in the recirculation line between the reservoir tank exit and the second membrane contactor. We stored the samples in 1.5-mL Eppendorf tubes at -20 °C until analyzing them in batches of approximately 60 units. We measured the concentrations of simple carboxylates and ethanol with a gas chromatograph, and succinate, lactate, pyroglutamate, and pyruvate with a high performance liquid chromatograph (Analytical procedures, ESI †) and normalized all given values to mmol carbon atoms (mmol C). Normalization simplifies the quantitative allocation of carbon atoms, especially to carboxylates of different chain lengths. We modified the calculation of the volumetric production rate from a previous report by compensating for the dilution of carboxylates in the alkaline extraction solution (Bioreactor Equations, ESI †).^23^ To trace isotope incorporations, we measured the ^13^C-isotopomer abundances of produced carboxylates with a gas chromatograph coupled to a mass spectrometer (Analytical procedures, ESI †). We measured non-labeled carboxylate fragmentation patterns as standards to determine which carbon-chain segments are analyzable (**Fig. S2-S3**, ESI †) and to compensate for naturally occurring isotopes (Mass spectrometry equations, ESI †). Fractional isotopomer ratios represent a weighted average between biological duplicates.

### *Clostridium kluyveri* isolation

We isolated a strain of the bacterial species *C. kluyveri* from the AF reactor, which shows deviating phenotypic traits from the type strain. For the isolation, we anaerobically sampled 1 mL of fermentation broth on Day 700, which we transferred into a sealed serum bottle. Under an anaerobic atmosphere, we spread 1 µL of the fermentation broth on an agar plate. The plate contained a modified reinforced clostridial medium with 15.78 g L^-1^ (342.5 mM) supplemented ethanol (*Clostridium kluyveri* isolate, ESI †). We added ethanol to give *C. kluyveri* a competitive advantage by creating a favorable selection pressure. After inoculating, we stored the agar plate at 30 °C in a gas-tight jar filled with a nitrogen/carbon dioxide gas mixture (80/20% v/v) and an additional ethanol atmosphere to prevent outgassing. After detecting microbial colonies through visual inspection, we picked them with a 1-mL syringe needle. We transferred the selected colonies separately into serum bottles. The serum bottles contained modified reinforced clostridial medium. Next, we incubated the serum bottles at 30 °C with shaking. After the growth phase, we mechanically lysed the cells and sequenced the 16S rRNA gene region to clarify the taxonomic classification. We verified the *C. kluyveri* isolate by sequencing of the 16S rRNA gene region and, subsequently, elucidated its product spectrum in bottle experiments (*Clostridium kluyveri* isolate, ESI †).

### Microbiome analysis

#### Metagenomics analysis

We gathered biomass samples from the fermentation broth of both bioreactors at four consecutive time points throughout the operating period. To collect biomass samples, we obtained 25-mL fermentation broth using a 50-mL syringe from two locations: **(1)** the bioreactor columns on Day 9 and 283 (both bioreactors), on Day 455 (only AF reactor) or Day 397 (only UASB reactor); and **(2)** the filter module on Day 701 (both bioreactors). The sampling procedure involved mixing the fermentation broth by quickly pulling and releasing the liquid with the 50-mL syringe ten times, effectively resuspending the immobilized biomass. Next, we physically lysed the contained microbes, extracted and prepared DNA, and used long-read sequencing DNA sequencing to obtain the metagenome of the microbial community (Metagenomics, ESI †). The results are shown in relative abundance, which we calculated by dividing the aligned base pairs allocated to a specific taxon, normalized by sample size, by the total number of aligned base pairs. We calculated the Pearson correlation coefficient to show correlations between relative species abundance and volumetric *n*-caprylate production rates. We reported the Pearson correlation coefficient as *r* (degrees of freedom) = Pearson *r, p* = *p* value. The critical value for positive correlation was *r* = .707 at a significance level of *p* < .05 and n = 8.

#### Comparative metaproteomics analysis

To detect proteins related to *n*-caprylate production, we compared mixed-liquor samples from both bioreactors during high and low volumetric *n*-caprylate production rates. We took the first sample on Day 408 during regular operating conditions. For the second sample, we intended to let the volumetric *n*-caprylate production rate of both bioreactors purposely collapse by adding L-cysteine into the medium. On Day 422, we collected the second sample. Later, we noticed that we did not suppress the *n*-caprylate productivity in the AF reactor as intended, so we did not show the related results.

The samples were 200 mL, each collected with a 50-mL syringe and distributed into 50-mL Falcon tubes. Immediately after, we centrifuged the Falcon tubes at 3450 ×*g* (4 °C) for 10 min and discarded the supernatants. We resuspended the emerged biomass pellets in 50 mM, pH 8 tris buffer containing 480 mg urea, transferred them into 2-mL Eppendorf tubes, centrifuged them again at 15000 ×*g* (at 4 °C), discarded the supernatants, and stored them at -20 °C until further use.^32^ We processed and analyzed the samples according to previous reports (Metaproteomics, ESI †).^33^ We did not include eukaryotes in the metaproteomics analysis.

#### Metabolomics analysis

We sampled 3 mL of fermentation broth from both bioreactors on Day 866 and 937 and aliquoted the samples into three Eppendorf tubes, resulting in 1 mL per sample. The AF reactor produced 1.8 mmol C L^-1^ d^-1^ of *n*-caprylate on Day 866 and 30.7 mmol C L^-1^ d^-1^ on Day 937. The UASB reactor produced 3.1 mmol C L^-1^ d^-1^ and 26.8 mmol C L^-1^ d^-1^ *n*-caprylate during the same days, respectively. We sampled twice per timepoint, once for extracellular metabolite analysis and again for combined extra- and intracellular metabolite analysis. We kept the samples on ice for subsequent steps to reduce metabolite degradation. We added 1 mL of quenching solution consisting of an acetonitrile methanol mixture (1:1 v/v; 4 °C) to lyse microbial cells in the combined intra- and extracellular metabolite samples. Afterward, we vortexed all samples for 10 s and centrifuged them at 9000 ×*g* and 4 °C for 5 min. We removed 100-µL supernatant of the extracellular and 200-µL supernatant of the combined intra- and extracellular samples and sterile-filtered (pore size 0.22 µM) it into a new Eppendorf tube (different volumina due to dilution with quenching solution). Finally, we evaporated the water and quenching solution from the samples by purging a steady nitrogen gas flow on the opened tubes. After evaporation, we stored the samples at -80 °C until further analysis (Metabolomics, ESI †).^34^

## Results

### We produced *n*-caprylate long-term and at relatively high rates during Period I

We operated an AF and UASB reactor independently with continuous feed and in-line product extraction. The medium contained ethanol and acetate as the sole substrate, and we omitted yeast extract. During the first 540 days (Period I), we observed low volumetric *n*-butyrate production rates and considerable volumetric *n*-caproate production rates (**Fig. 1**), while *n*-caprylate was the main product in both bioreactors (**Fig. S4B,D**, ESI †). The AF reactor outperformed the UASB reactor, producing a maximum of 190.0 ± 8.4 mmol C L^-1^ d^-1^ (0.14 g L^-1^ h^-1^) *n*-caprylate (**Fig. 1**). At peak production levels, the *n*-caprylate selectivity was 69 ± 5.7% throughout an operating period of 10 days (Day 142-152; **Fig. 1A**). The UASB reactor had a lower maximum volumetric *n*-caprylate production rate at 131.1 ± 4.7 mmol C L^-1^ d^-1^ and a 52.4 ± 10.5% selectivity for *n*-caprylate throughout 10 days (Day 221-231; **Fig. 1B**). During the peak performance, neither ethanol nor carboxylates accumulated in the fermentation broth (no overfeeding) (**Fig. 1, Fig. S4A,C**, ESI †). For both bioreactors, the volumetric *n*-caprylate production rates were consistent during Period I outside of the treatments (2xA-C in **Fig. 1**). However, they fluctuated the most among the measured carboxylates (46% standard deviation during Period I in the AF reactor and 36% in the UASB reactor) (**Fig. 1**).

**Fig. 1.**
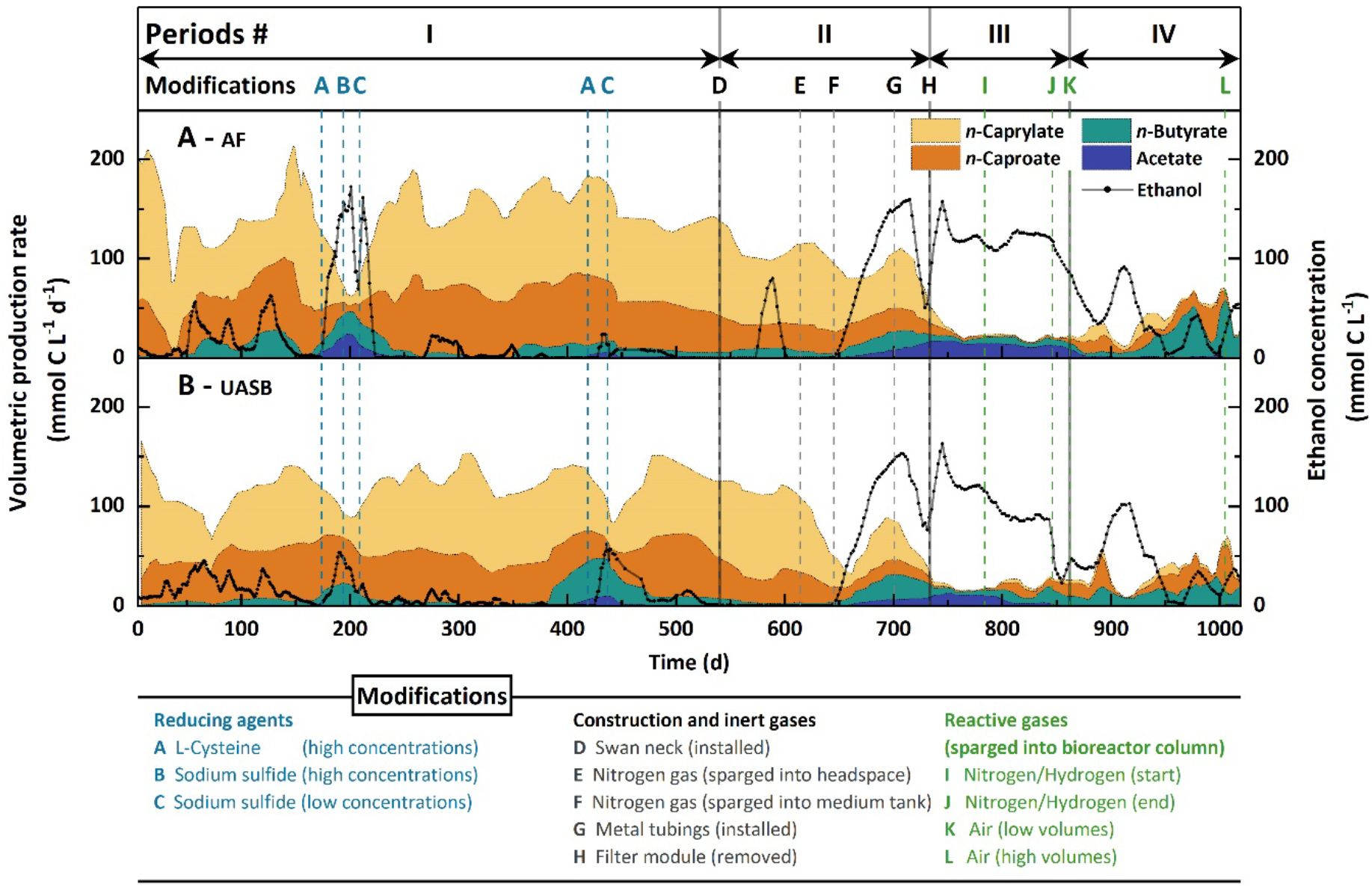
Volumetric production rates of carboxylates, ethanol concentrations, and modifications to the **A**. AF reactor and **B**. UASB reactor during the study. The volumetric production rates were calculated as a six-measurement moving average. The volumetric production rates and the ethanol concentration values were determined from samples that were collected daily or every other day. The operating Periods I – IV and modifications to the bioreactors are indicated and described in the experimental section. The modification timelines mark the first experimental day of implementation. All the following days include the respective modification except when indicated otherwise (*i.e*., reducing agent dosing and reactive gases were added alternatingly).

We added 0.3 g L^-1^ L-cysteine (within the range of general Clostridia nutrient media) to the feed medium of the AF and UASB reactors during Day 174-194 and Day 419-449, which led to sharp drops in volumetric *n*-caprylate production rates (2xA in **Fig. 1**).^35^ Two times, the addition of L-cysteine to the medium led to a breakdown in the volumetric *n*-caprylate production rates (Day 174-194 in the AF reactor: 13-fold decrease; Day 419-449 in the UASB reactor: 8-fold decrease). The other two times, we observed milder decreases in the volumetric *n*-caprylate production rates (Day 419-449 in the AF reactor and Day 174-194 in the UASB reactor). During the breakdown periods, the ethanol, acetate, and *n*-butyrate concentrations in the fermentation broths of both bioreactors increased drastically (2xA-C in **Fig. 1** and **Fig. S4A,C**, ESI †). Changing the reducing agent to a low dose of sodium sulfide (0.02 g L^-1^ d^-1^) restored the high volumetric *n*-caprylate production rates in all instances (2xC in **Fig. 1**). The breakdowns of *n*-caprylate production after the addition of reducing agent led to our hypothesis that oxygen played a fundamental role in the process.

### The availability of oxygen was critical to the performance of both bioreactors

During Period I, oxygen intruded into the bioreactors through diffusive processes. To close the systems and cut off all oxygen sources during Period II, we installed a swan-neck (D in **Fig. 1**), sparged the headspace and medium tank with a continuous nitrogen gas flow (E and F in **Fig. 1**), and changed the plastic tubing to metal tubing (G in **Fig. 1** and **Table S2**, ESI †). We measured dissolved oxygen concentrations in the fermentation broth in response to these system modifications (**Table S3**, ESI †). The extraction systems for extracting medium-chain carboxylates additionally removed oxygen from the recirculating fermentation broth (Positions 4+5, Explanatory figure for **Table S3**, ESI †). We measured the same decrease in oxygen concentrations in the extraction systems for the AF and UASB reactors: upstream 0.09 ± 0.01 mg O_2_ L^-1^; and downstream 0.04 ± 0.01 mg O_2_ L^-1^. On the other hand, the filter modules increased the oxygen concentration in the recirculating fermentation broth (Positions 6+7, explanatory figure for **Table S3**, ESI †). We measured oxygen concentrations upstream and downstream of the filter module of 0.19 ± 0.1 mg O_2_ L^-1^ and 0.48 ± 0.07 mg O_2_ L^-1^ for the AF reactor. For the UASB reactor, the oxygen concentrations were 0.17 ± 0.02 mg O_2_ L^-1^ and 0.42 ± 0.02 mg O_2_ L^-1^, respectively. Therefore, for the start of Period III, we removed the filter modules from the bioreactors to further circumvent oxygen intrusion (H in **Fig. 1**).

While single modifications had variable effects on the volumetric *n*-caprylate production rates of the bioreactors (**Table S2**, ESI †), cumulatively, they had an inhibitory impact. Between Periods I to III, the volumetric *n*-caprylate production rates decreased from 93.0 ± 43.1 mmol C L^-1^ d^-1^ to 1.2 ± 2.1 mmol C L^-1^ d^-1^ in the AF reactor and from 73.5 ± 26.7 mmol C L^-1^ d^-1^ to 2.5 ± 3.3 mmol C L^-1^ d^-1^ in the UASB reactor (**Fig. 1** and **Table S2**, ESI †). Also, the product spectrum shifted toward higher volumetric *n*-butyrate production rates of up to 33.2 ± 4.8 mmol C L^-1^ d^-1^ for the AF reactor and 44.5 ± 5.0 mmol C L^-1^ d^-1^ for the UASB reactor (**Fig. 1**). Lowering the total organic loading rate stepwise to 80% and 60% and sparging the bioreactor column with a nitrogen/hydrogen gas mixture had minor relieving effects on the residual ethanol concentrations (between I and J in **Fig. 1**). During this time, the UASB reactor started producing small amounts of *n*-caprylate at up to 11.6 ± 4.5 mmol C L^-1^ d^-1^ (**Fig. 1**). Interestingly, the combined effect of stopping the nitrogen/hydrogen gas supply and a decreased total organic loading rate toward the end of Period III resulted in a substantial decrease in ethanol concentrations (J in **Fig. 1**). Thus, the nitrogen/hydrogen gas supply had prevented a performance improvement, possibly due to flushing out oxygen.

During Period IV, we studied the effect of introducing low volumes of air (∼21% oxygen) through the bottom of the bioreactor vessels (K in **Fig. 1**). Apart from a production-rate drop with an accompanied ethanol concentration spike due to an operating error between Day 899 and 929, the residual ethanol concentration in the fermentation broths of both bioreactors decreased further to zero on Day 958 without acetate accumulation (**Fig. 1**). There was a clear improvement in the volumetric *n*-caproate and *n*-caprylate production rates in both bioreactors (**Fig. 1**), wherein the volumetric *n*-caprylate production rate peaked at 38.8 ± 5.6 mmol C L^-1^ d^-1^ in the AF reactor, albeit this rate remained inconsistent (**Fig. 1A**). An increase in the supplied air volume toward the end of the experiment led to near zero volumetric *n*-caproate and *n*-caprylate production rates in both bioreactors (L in **Fig. 1**), leading us to stop the operation on Day 1019. The relatively high selectivity of *n*-butyrate after Day 958 indicated that we had not applied oxygen optimally to restore *n*-caprylate production rates compared to Period I for both bioreactors (**Fig. 1**). We over-applied oxygen, which can be improved by employing a more finely adjustable oxygen application system or separation of the aerobic and anaerobic phases.

### Bottle experiments confirmed the enhanced *n*-caprylate production rate in response to oxygen supply

Following up on the results from the bioreactor operation, we investigated the effect of oxygen availability on carboxylate production with bottle experiments utilizing a modified medium containing [1,2-^13^C_2_]-labeled ethanol and no acetate (**Fig. 2**). Fresh fermentation broth from the AF reactor served as the inoculum, containing the pertinent carboxylate-producing microbial community with undefined intra- and extracellular metabolites. On Day 0 of the experimental period, we inoculated with and without exposure to oxygen (*i.e*., henceforth termed aerobic and anaerobic treatment). The beginning phase exhibited a pronounced contrast in pH and cell density progression between the two treatments (**Fig. S5A-B**, ESI †). For the aerobic treatment, the pH rose from 5.0 to 6.9 ± 0.1, while the OD_600_ increased nearly 14-fold from 0.1 ± 0.1 to 1.36 ± 0.1 during the first four days. After Day 4 of the experimental period, the pH in both bottles decreased to around 5.0, which is close to the pKa (∼4.8–4.9) of the produced carboxylic acids (**Fig. S5A**, ESI †). In contrast, for the anaerobic treatment, the pH remained acidic throughout the experiment, while the OD_600_ did not exceed 0.35 (**Fig. S5A-B**, ESI †).

**Fig. 2.**
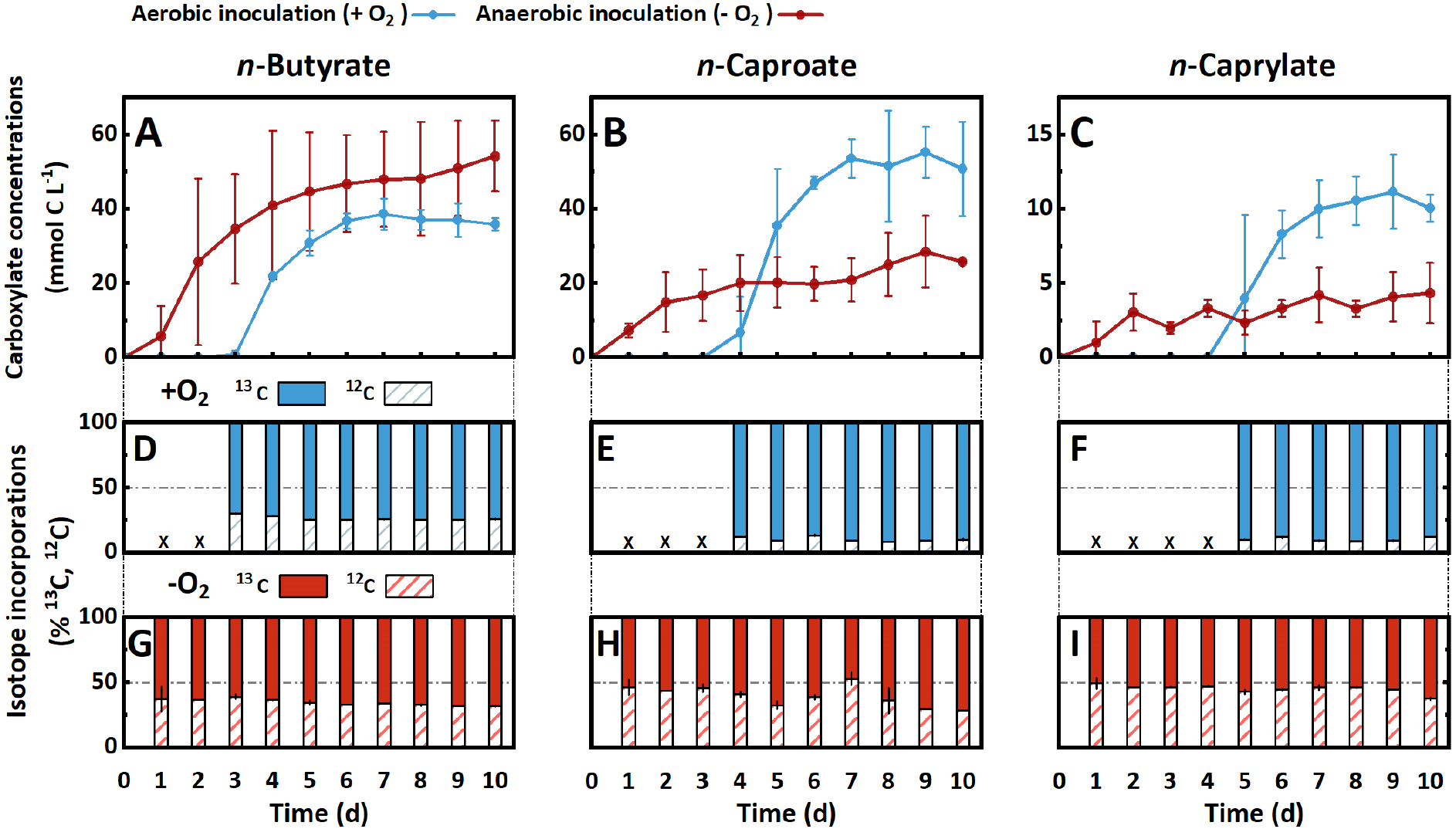
**A-C**. Carboxylate concentrations and **D-I**. total incorporation percentages of ^13^C and ^12^C isotopes of *n*-butyrate (left column), *n*-caproate (middle column), and *n*-caprylate (right column) throughout the 10-days bottle experiments. The aerobically inoculated bottles are light blue, and the anaerobically inoculated bottles are shown in red. The aerobic treatment produced *n*-caproate and *n*-caprylate with an onset delay at higher concentrations and more rapidly. We calculated the percentual isotope incorporations using **Eq. S8-10** and **Eq. S11-12** (ESI †). A more thorough incorporation of ^13^C-ethanol was observed in the aerobic treatment (middle and right bottom row). Error bars represent the standard error between biological duplicates.

The product spectrum of the aerobic and anaerobic treatments comprised *n*-butyrate, *n*-caproate, and *n*-caprylate (**Fig. 2A-C**). The aerobic treatment had a delayed onset of rapid carboxylate production followed by a plateau in carboxylate concentrations (**Fig. 2A-C**). Successively, we measured *n*-butyrate after three days, *n*-caproate after four days, and *n*-caprylate after five days (**Fig. 2A-C**). For the anaerobic treatment, the three carboxylates were present in measurable concentrations from Day 1, with a steady increase of the *n*-butyrate concentration to 54.2 ± 9.6 mmol C L^-1^ whereas the *n*-caproate and *n*-caprylate concentrations increased sluggishly to 25.8 ± 0.2 and 4.3 ± 2.0 mmol C L^-1^, respectively (**Fig. 2A-C**). Notably, compared to the anaerobic treatment, the aerobic treatment resulted in: **(1)** a lower maximum *n*-butyrate concentration (35.9 ± 1.7 mmol C L^-1^) (**Fig. 2A**); **(2)** two-fold higher *n*-caproate and *n*-caprylate concentrations (50.8 ± 12.6 and 10.1 ± 0.9 mmol C L^-1^, respectively) (**Fig. 2B-C**); **(3)** a higher consumption of ethanol (106 ± 19.8 *vs*. 66 ± 8.5 mmol C L^-1^, respectively) (**Fig. S5C**, ESI †); and **(4)** more total carbon in the carboxylate product (96.7 ± 15.3 *vs*. 84.3 ± 7.3 mmol C L^-1^, respectively) (**Fig. S5D**, ESI †). Finally, the yield (% mol/mol, carboxylates produced/ethanol consumed) was lower in the aerobic treatment (91.3 ± 24.5%) compared to the anaerobic treatment (127.8 ± 15.5%). The higher than 100% ethanol-based yield for the anaerobic treatment implied that the anaerobic treatment produced more carboxylates than it consumed ethanol. Because ethanol was the only carbon source in the modified medium, utilizable metabolites had to be introduced as part of the initial inoculum.

### Aerobic and anaerobic treatments for bottle experiments displayed ^13^C-labeling pattern discrepancies

Monitoring the flow of ^13^C-labeled ethanol into carboxylates revealed further distinctions between the two treatments. The incorporation of ^13^C atoms was higher for the aerobic treatment than for the anaerobic treatment (**Fig. 2D-I**), which confirmed the ethanol-based yield results above (a lower yield for the aerobic treatment implies lower utilization of other metabolites compared to ethanol). The higher maximum labeling extent in the aerobic treatment than in the anaerobic treatment (91% *vs*. 71%) indicated that labeled ethanol was the primary carbon atom source for carboxylate production (**Fig. 2D-I**). However, for the anaerobic treatment, the labeling pattern implies the utilization of a combination of labeled ethanol and an unlabeled metabolite for the production of carboxylates. Between the two treatments, we observed a reversed labeling extent of *n*-butyrate, *n*-caproate, and *n*-caprylate. On the one hand, for the aerobic treatment, *n*-butyrate had the lowest average labeling fraction (73.5 ± 0.3%), while *n*-caproate and *n*-caprylate were labeled at higher fractions (89.7 ± 0.4% and 89.6 ± 0.5%, respectively) (**Fig. 2D-F**). On the other hand, for the anaerobic treatment, *n*-butyrate had the highest average labeled fraction (65.3 ± 1.5%), while *n*-caprylate had the lowest fraction (54.9 ± 1.2%) (**Fig. 2G-I**).

We also observed that the labeled fraction did not change substantially for the aerobic treatment (**Fig. 2D-F**). For the anaerobic treatment, however, the labeled fraction increased throughout the experimental period (**Fig. 2G-I**). For example, the labeled fraction of *n*-caproate increased from 53.6 ± 6.0% on Day 1 to 71.2 ± 0.1% on Day 10 (**Fig. 2H**). This stale and increased labeling shows that for the aerobic treatment, the stoichiometries of the metabolic reactions were determining the labeling extends of medium-chain carboxylates. Meanwhile, for the anaerobic treatment, there appeared to be a depletion in the unlabeled metabolite from the fermentation broth after some time.

We expanded the monitoring of isotope incorporations into carboxylates by calculating the fractional isotopomer abundances per analyzable carbon-chain segment to ascertain labeling patterns (**Fig. S2** and **Eq. S8-10**, ESI †). Counting carbon atoms on the carboxylate carbon chain always starts from the carbon in position 1, which is part of the carboxyl group. Thus, the two-carbon-chain segment is composed of the carbon atoms in positions 1-2, and the three-carbon-chain segment of the carbon atoms in positions 1-3, etc. (**Fig. S2**, ESI †), until the maximum eight-carbon-chain segment in *n*-caprylate. Incorporation events to the carbon chain of carboxylates (*i.e*., chain elongation) occur at position 1 in the carboxyl group, which is the functional group.

The carboxylates in the aerobic treatment tended to contain fully or close-to-fully labeled carboxylate isotopomers (**Fig. 3** and **Fig. S6-S8**, ESI †), as already reflected by the total ^13^C-labeled fractions (**Fig. 2**). For instance, 75% of the six-carbon-chain segments (positions 1-6) of *n*-caprylate contained six ^13^C atoms throughout the experimental period (red bars in **Fig. 3C**). In contrast, for the anaerobic treatment, there was a diverse distribution of isotopomer abundances for the six-carbon-chain segment for *n*-caprylate (**Fig. 3D**). In addition, for the aerobic treatment, we observed differences in the labeling patterns of different carboxylates where there was a higher fully labeled two-carbon-chain segment (*m/z* 60-62, M+2) fraction (88.1 ± 3.7%) for *n*-butyrate than for *n*-caproate and *n*-caprylate (79.3 ± 1.7% and 80.8 ± 0.6%, respectively) (**Fig. S6A, Fig. S7A**, and **Fig. S8A**, ESI †). Because the two-carbon-chain segment shows the most recent incorporation of carbon atoms into the carboxylate, this higher labeling of the two-carbon-chain segment shows that labeled ethanol preferably flowed into the recent addition to form *n*-butryate for the aerobic treatment.

**Fig. 3.**
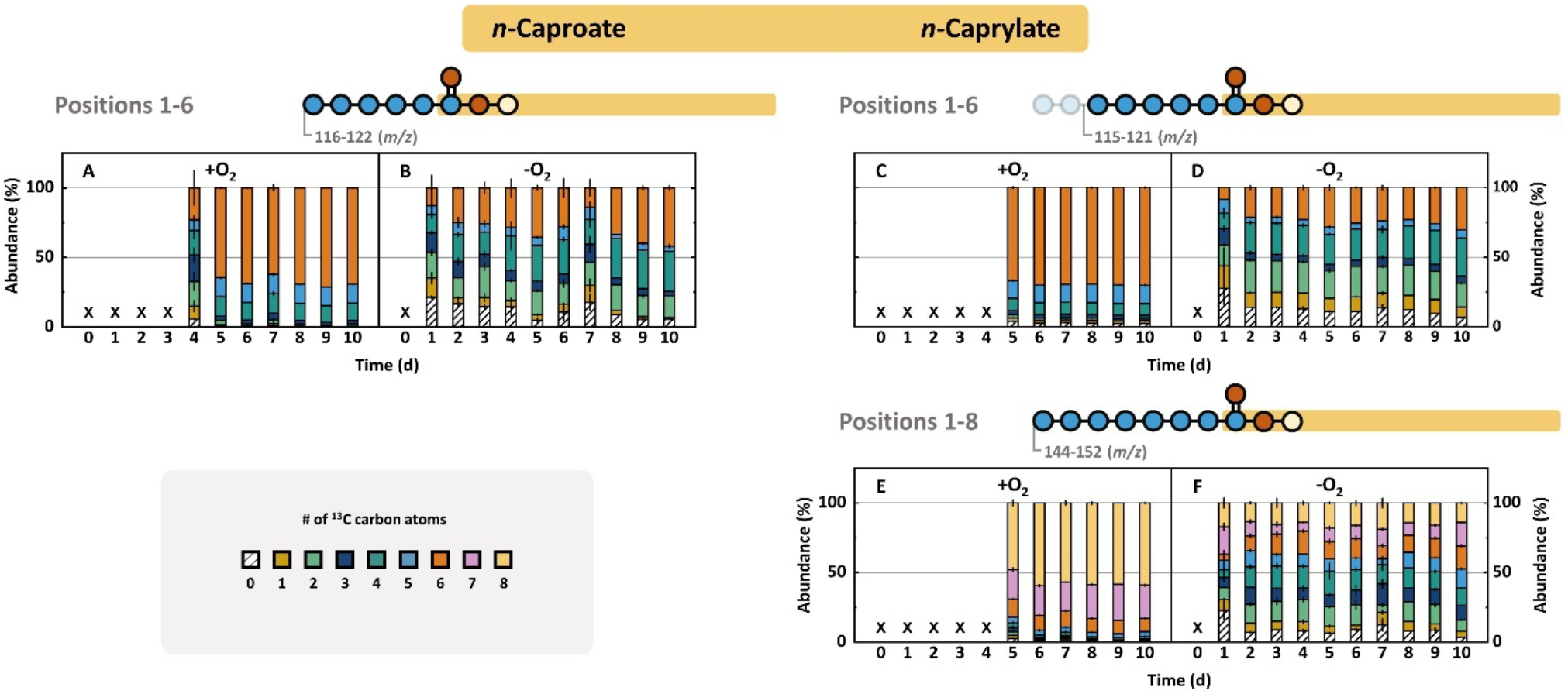
Fractional isotopomer abundances of *n*-caproate and *n*-caprylate throughout the 10-day experimental period. The isotopomer distribution of the complete *n*-caproate compound in the **A**. aerobic (+O_2_) or **B**. anaerobic (-O_2_) treatment. The isotopomer distribution for the six-carbon-chain segment (positions 1-6) of *n*-caprylate in the **C**. aerobic (+O_2_) or **D**. anaerobic (-O_2_) treatment. The isotopomer distribution for the whole *n*-caprylate compound (positions 1-8) in the **E**. aerobic (+O_2_) or **F**. anaerobic (-O_2_) treatment. The isotopomers are arranged from bottom to top by increasing ^13^C-share. The data is read as follows: A 25% abundance of the isotopomer with two ^13^C atoms in the six-carbon-chain of *n*-caprylate means that, on average, one in four *n*-caprylate molecules contains two heavy isotopes in positions 1-6. The positions of those ^13^C atoms are not specified. The aerobic treatment contained carboxylates with uneven distribution and a trend toward fully labeled carbon species. The anaerobic treatment contained well-distributed isotopomer abundances with a preference for even-numbered integration in the six-carbon-chain segments. The depicted values derive from daily sampling, and the errors represent the standard error between biological duplicates.

For a given carboxylate, we cannot *directly* infer the position of the isotopes on the carbon chain. However, our gas chromatograph mass spectroscopy method includes hard ionization after carboxylate separation by the column, generating each carbon-chain segment for the given carboxylate (**Fig. S2**, ESI †). With this information, we traced the incorporation steps of ^13^C atoms into carboxylates by *indirectly* inferring the position of the isotopes on the carbon chain by comparing carbon-chain segments (*e.g*., by comparing the two-carbon-chain segment to the three-carbon-chain segment of given carboxylates) from lower-to-higher position number (reverse chronology of ^13^C-incorporation). It is scientifically proven with *C. kluyveri* that chain elongation from acetate to *n*-butyrate (1^st^ chain-elongation step) and from *n*-butyrate to *n*-caproate (2^nd^ step) occurs with two carbon atoms from acetyl-coenzyme A (acetyl-CoA) at the same time at position 1, which we refer to as the current model (**Fig. 4A**). ^36^ However, this model has not yet been scientifically proven for chain elongation from *n*-caproate to *n*-caprylate (3^rd^ step). Therefore, we retraced the incorporation steps of carbon atoms all the way from acetate to *n*-caprylate in the three elongation steps for even-chain carboxylates (**Fig. 3** and **Fig. S6-S8**).

**Fig. 4.**
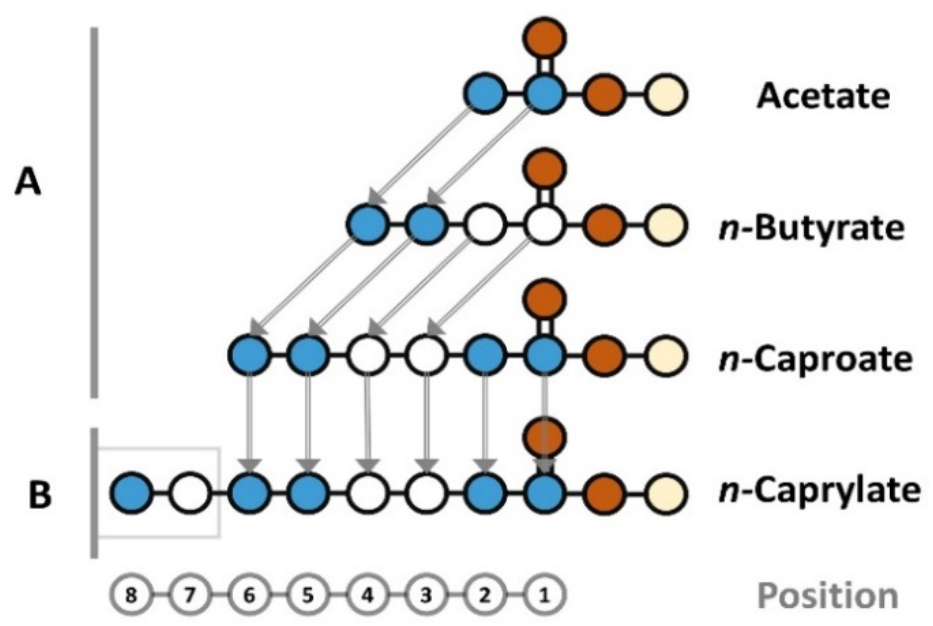
Proposed molecular assembly of observed carboxylates in our study. The position on the carbon chain is shown shown at the bottom for reference. The addition of carbon atoms into carboxylates occured with two carbon atoms at a time in all cases because the concentrations of odd-numbered carboxylates was below the detection limit. **A**. The elongations from acetate to *n*-butyrate and from *n*-butyrate to *n*-caproate proceeded as similarly labeled, two-carbon-atom integrations at position 1. **B**. The elongation from *n*-caproate to *n*-caprylate proceeded with differently labeled, two-carbon-atom integration at position 7 (**Fig. 3D,F**).

For the anaerobic treatment, the two-carbon-chain segments of *n*-butyrate, *n*-caproate, and *n*-caprylate displayed a similar labeling preference for: **(1)** two ^13^C-isotopes (M+2, fully labeled) at an 73.9 ± 6.3% average abundance; **(2)** one ^13^C-isotope (M+1) at a negligible fraction at an 3.4 ± 1.6% average abundance; and **(3)** zero ^13^C-isotopes (M+0) at a 21.6 ± 3.3% average abundance (**Fig. S6B, Fig. S7B**, and **Fig. S8B**, ESI †). This result indicates that for our experiment: **(i)** both carbon atoms of acetyl-CoA would be equally labeled (either with ^13^C-labeled atoms from fully labeled ethanol or non-labeled carbon atoms from a metabolite in the fermentation broth); and **(ii)** the recent integration step occurs with at least two carbon atoms at a time rather than just with one carbon atom. To exclude that the incorporation step does not involve more than two carbon atoms at position 1, we compared the two-carbon-chain segments to the three-carbon-chain segments of carboxylates. If the current model is true, then a three-carbon-chain segment would contain an earlier incorporation in position 3 with positions 1 and 2 from the recent incorporation. Thus, position 3 could be differently labeled than the equally labeled positions 1 and 2 (**Fig. 4A**).

To investigate whether there is a break in the incorporation mechanism between position 2 and 3, which would be caused by a two-carbon-step addition, we first compared the fully labeled fraction of the two-carbon-chain segment (*m/z* 60-62; M+2) to the fully labeled fraction of the three-carbon-chain segment (*m/z* 73-76; M+3). Indeed, for the anaerobic treatment, the two-carbon-chain segments had higher abundances of fully labeled fractions than the three-carbon-chain segments: 76.4 ± 4.5% *vs*. 55.6 ± 3.7% for *n*-butyrate, 72.7 ± 2.9% *vs*. 49.0 ± 2.5% for *n*-caproate, and 72.7 ± 1.4% *vs*. 50.4 ± 1.9% for *n*-caprylate throughout the experimental period (**Fig. S6D, Fig. S7D**, and **Fig. S8D**, ESI †). In addition, we observed one ^13^C-labeled isotopomers in the three-carbon-chain segment for the anaerobic treatment with appreciable average abundances of 10.6 ± 0.8% for *n*-butyrate, 15.0 ± 2.7% for *n*-caproate, and 11.8 ± 0.5% for *n*-caprylate throughout the experimental period (**Fig. S6D, Fig. S7D**, and **Fig. S8D**, ESI †). For our experiment, the one ^13^C-labeled isotope would be located at position 3 while position 1 and 2 would be mostly unlabeled (**Fig. 4**). We observed approximately twice the average abundance for two ^13^C-labeled isotopomers in the three-carbon-chain segment compared to the one ^13^C-labeled isotopes with 18.4 ± 0.4% for *n*-butyrate, 22.5 ± 1.0% for *n*-caproate, and 24.4 ± 0.9% for *n*-caprylate (**Fig. S6D, Fig. S7D**, and **Fig. S8D**, ESI †). For our experiment, this identified mostly an unlabeled position 3, while positions 1 and 2 were labeled.

Considering the isotopomer distributions in the two and three-carbon-chain segments, we conclude that the recent carbon addition step in *n*-butyrate, *n*-caproate, and *n*-caprylate occurred with two carbon atoms at a time and not three or more. The higher abundance of the fully labeled two-carbon-chain segment (M+2) than the fully labeled three-carbon-chain segment (M+3) indicated a break in the carbon incorporation mechanism. The emergence of the one labeled carbon atom (M+1) in the three-carbon-chain segment revealed the earlier carbon addition at positions 3 and 4, which was uncoupled from the recent addition in positions 1 and 2.

This two-carbon *recent* addition conclusion would only hold when it occurred at position 1 according to the current model, and this was not scientifically proven to occur for *n*-caprylate. We observed that the labeling pattern of the complete *n*-caproate compound with six-carbon atoms matched the labeling pattern of the six-carbon-chain segment (positions 1 and 6) of *n*-caprylate for the anaerobic treatment (**Fig. 3B,D**). Clearly, *n*-caproate is the substrate for *n*-caprylate. However, such a close match of the six-carbon-chain segments for both *n*-caproate and *n*-caprylate could mean two possibilities: **(1)** a very similar labeling pattern for the two-carbon atoms for the recent addition at position 1 for both *n*-caproate and *n*-caprylate; or **(2)** labeling at the other side of *n*-caproate from position 7, while the six-carbon-chain segment for *n*-caprylate remains the same when compared to *n*-caproate (**Fig. 4B**).

The latter possibility can be checked by comparing positions 7 and 8 in *n*-caprylate with positions 5 and 6 in *n*-caproate, which should have the same labeling pattern according to the current model (**Fig. 4**). Here, we found a discrepancy. For the aerobic treatment, we found a large share of the seven-labeled isotopomer (M+7) at 22.8 ± 2.1% in the eight-carbon-chain segment of *n*-caprylate (**Fig. 3E**). Similarly, for the anaerobic treatment, we observed the current model for the addition of the two-carbon atoms at position 1 to continue until the six-carbon-chain segment of *n*-caprylate, verifying the scientifically proven chain elongation for *n*-caproate (**Fig. S8B, Fig. S8D**, and **Fig. 3D**, ESI †). The current model fell apart for the chain elongation from the *n*-caproate into the *n*-caprylate molecule (**Fig. 3F**), which would imply a different type of molecular assembly in position 7 instead of the integration at position 1 (**Fig. 4B**). Still, the integration would have occurred by two-carbon atoms at a time, but now each carbon was labeled differently, albeit the pathway is unknown to us.

### The microbial community encompassed all three domains of life in distinct ecological niches

We used long-read metagenome sequencing to assess the microbial communities of both long-term operated bioreactors (*i.e*., AF and UASB) at three timepoints for the bioreactor vessels during Period I (at Day 9, Day 283, and Day 397 or 455) and on time point for the filter moduls toward the end of Period II (at Day 701) (**Fig. 5**). For the metagenomics samples, we captured 462 distinct operational taxonomic units in the AF reactor and 317 in the UASB reactor. The microbial community in the AF reactor had a higher species richness and was more stable, evidenced by the averaged Shannon diversity index of 2.46 ± 0.12 compared to 1.88 ± 0.31 in the UASB reactor. The metagenome data showed low relative abundances of eukaryotic fungi and high relative abundances of bacteria and methanogenic archaea (**Fig. 5, Fig. S1C**, ESI †).

**Fig. 5.**
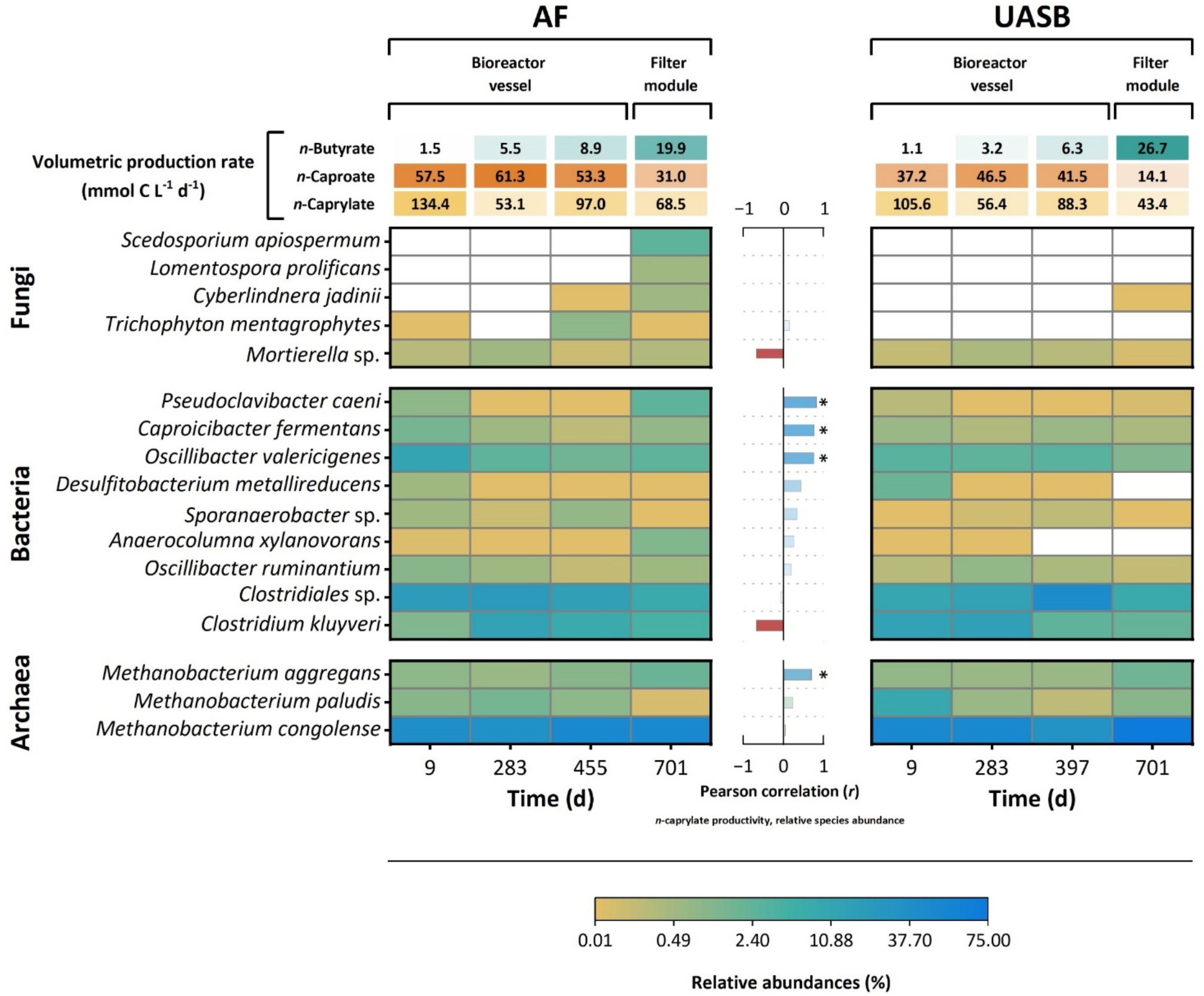
Heat map of relative microbial species abundances of the core community and volumetric carboxylate production rates in four metagenomics samples from the AF and UASB reactors during operating Periods I and II. The volumetric production rates of *n*-butyrate, *n*-caproate, and *n*-caprylate of the bioreactors during the displayed sampling days is shown on the top. We collected the first three samples from the bioreactor vessels and the last samples from the filter modules of each bioreactor. The cutoff value was a relative abundance of 0.5% for fungi and 1% for bacteria and archaea in at least one of the metagenome samples. We ordered the microbial species according to their domain into Eukaryota (comprising members of the kingdom fungi), bacteria, and archaea. Species in each domain were ordered hierarchically from top to bottom, from highest correlation to the volumetric *n*-caprylate production rate to lowest. The Pearson correlation values for the volumetric *n*-caprylate production rate and relative species abundance are displayed as a bar graph between the heatmaps. Only samples from the bioreactor vessels were considered in the correlation calculation: Light blue values indicate a positive correlation, white values do not correlate, and red values correlate negatively. We highlighted critical values with an asterisk. *Scedosporium apiospermum* and *Lomentospora prolificans* were found exclusively in one sample. Therefore, the correlation values could not be determined. Fields with relative abundances below 0.01% are shown in white color. *P. caeni*, which is an aerobic bacterium of the Microbacteriacaea family, showed the highest correlation to *n*-caprylate production and was most abundant in the filter module of the AF reactor. The color assignment of the relative species abundance is on a logarithmic scale.

To represent the core community, we selected fungal species with at least 0.5% and bacterial and archaeal species with at least 1.0% relative abundance in at least one of the metagenomics samples (**Fig. 5**). This core community, which accounted for 80.7% to 91.6% of relative abundance, consisted of five fungal, nine bacterial, and three archaeal species. The dominant community consisted of *Methanobacterium congolense* (33.7% to 74.2%), Clostridiales sp. (8.7% to 43.4%), *C. kluyveri* (1.4% to 17.3%), and *Oscillibacter valericigenes* (1.4% to 12.3%) (**Fig. 5**). The matches for Clostridiales sp. consisted of a mixture of uncharacterized species that did not match taxons on family, genus, or species level. Among others, the order Clostridiales comprises the families Clostidiaceae, Eubacteriaceae, Oscillospiraceae, and Lachnospiraceae, which include chain-elongating species.

Within the core community, most species are anaerobic (*i.e*., they can only tolerate minor amounts of oxygen). Only the fungal species are facultative anaerobic and the bacterium *Pseudoclavibacter caeni* is strictly aerobic (*i.e*., they all can thrive in the presence of oxygen). We found the species that thrive in the presence of oxygen either exclusively or at comparably higher relative abundances: **(1)** in the AF reactor; and **(2)** in the filter modules (**Fig. 5**). The relative abundance of *P. caeni*, which is a member of the Microbacteriaceae family, correlated significantly positively to the volumetric *n*-caprylate production rate [*r*(6) = .82, *p* = .01] (**Fig. 5**). Other species that correlated significantly positively to the volumetric *n*-caprylate production rate were *Caproicibacter fermentans* [*r*(6) = .77, *p* < .01], *O. valericigenes* [*r*(6) = .76, *p* < .01], and *Methanobacter aggregans* [*r*(6) = .71, *p* < .01] (**Fig. 5**). Both, *C. fermentans* and *O. valericigenes* are chain-elongating members of the Oscillospiraceae family. Similar to the other two highly abundant archaea (*Methanobacterium paludis* and *M. congolense*), *M. aggregans* is a hydrogenotrophic methanogen. The slight negative correlation of the model microbe for chain elongation, which is *C. kluyveri*, with the volumetric *n*-caprylate production rate [*r*(6) = -.68, *p* < .01], was noteworthy (**Fig. 5**).

We isolated the *C. kluyveri* strain from the AF reactor, and, as anticipated, bottle experiments with the isolated *C. kluyveri* species showed an absence of *n*-caprylate production (**Fig. S9**, ESI †). Moreover, the isolate did not produce *n*-caproate, which differs from the type strain *C. kluyveri* DSM 555. The *C. kluyveri* strain from our bioreactor showed 99.23% 16S rRNA gene similarity to DSM 555.^37^ Therefore, while our isolate strain was not a discrete operational taxonomic unit, it exhibited particular phenotypic features. The only carboxylate produced by our isolate was *n*-butyrate, demonstrating adaptive integration into the bioreactor microbiome.

### Tricarboxylic acid cycle-related proteins and metabolites were most positively related to *n*-caprylate production

To determine any correlations between specific protein expression and *n-*caprylate production rates for the long-term operated bioreactors, we compared a high and a low volumetric *n*-caprylate production rate treatment. Proteins belonging to the following metabolic pathways were more abundant in the high *n*-caprylate production rate treatment: the tricarboxylic acid (TCA) cycle (46% more); chain-altering pathways for carboxylates, such as the reverse-ß oxidation (34% more) and fatty acid degradation (21% more) (**Fig. 6A**) The reverse ß-oxidation pathway seemed the most active amongst these pathways, based on total protein abundance (**Fig. 6A**).

**Fig. 6.**
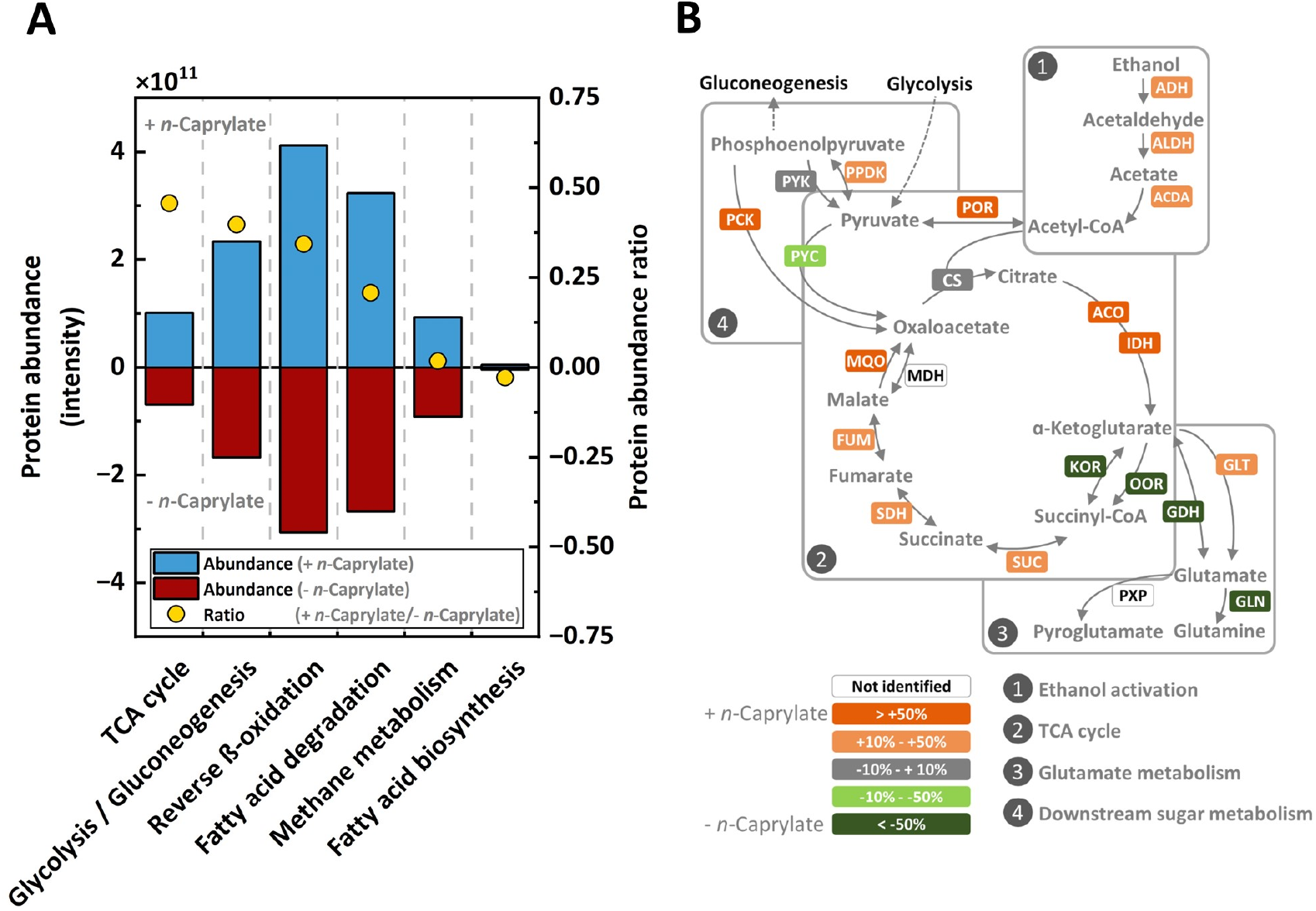
Metaproteomics profiling between treatments. Metaproteomics samples were obtained during a phase of a high volumetric *n*-caprylate production rate (+ *n*-Caprylate) and a low rate (-*n*-Caprylate), which was caused by L-cysteine addition (treatment). **A**. Total protein abundances and protein ratios between treatments mapped on central carbon metabolism, carbon chain-altering pathways of carboxylates, and methane metabolism. A ratio of 0.34, which we observed for reverse ß-oxidation as an abundant metabolic pathway, amounts to a 34% increase in total abundance for the high volumetric *n*-caprylate production-rate phase. The only negative ratio was observed for the fatty acid biosynthesis, and thus this pathway was negatively linked to the volumetric *n*-caprylate production rate. **B**. Percentage changes of proteins catalyzing the proposed aerobic preconversions. Proteins that correlate positively with volumetric *n*-caprylate production rates are highlighted in orange and proteins that correlate negatively in green. Protein abbreviations: alcohol dehydrogenase, ADH; aldehyde dehydrogenase, ALDH; acetate-CoA ligase, ACDA; pyruvate ferredoxin oxidoreductase, POR; pyruvate carboxylase, PYC; pyruvate orthophosphate dikinase, PPDK; pyruvate kinase, PYK; phosphoenolpyruvate carboxykinase, PCK; citrate synthase, CS; malate dehydrogenase, MQO; fumarate hydratase, FUM; succinate dehydrogenase, SDH; succinyl-CoA synthetase, SUC; 2-oxoglutarate ferredoxin oxidoreductase, KOR; 2-oxoacid ferredoxin oxidoreductase, OOR; aconitate hydratase, ACO; isocitrate dehydrogenase, IDH; glutamate dehydrogenase, GDH; glutamate synthase, GLT; glutamine synthetase, GLN; 5-oxoprolinase, PXP.

The only carbon chain-altering pathway of carboxylates unrelated to *n*-caprylate production was the fatty acid biosynthesis pathway with low total protein abundances, precluding it from *n*-caprylate production (**Fig. 6A**).

Furthermore, the abundance of proteins associated with the glycolysis and gluconeogenesis pathway was 40% higher in the high *n*-caprylate production-rate treatment than in the low *n*-caprylate production-rate treatment (**Fig. 6A**). Additionally, in contrast to the very high relative abundances of methanogenic archaea amongst the identified species (**Fig. 5**) we observed low abundances of the proteins involved in the pathway for methane production (**Fig. 6A**).

When combining the metagenomics and comparative metaproteomics data, we found that the dominating community and significantly positively correlating species expressed unique proteins during the high *n*-caprylate production treatment. We found two alcohol dehydrogenases (adh2, adhP) of *P. caeni* only present in the high *n*-caprylate production-rate treatment, suggesting efficient ethanol conversion under aerobic conditions by this species (**Table S4**, ESI †). Our results substantiate efficient ethanol conversion in the presence of oxygen, as proteins associated with oxygen stress were more abundant in the same treatment. (**Fig. S10**, ESI †). Additionally, we found unique proteins related to glutamate metabolism, as, for example, the phosphoserine aminotransferase (serC) of *O. valericigenes*, which was abundant for the high *n*-caprylate production-rate treatment (**Table S4**, ESI †).

We examined the TCA cycle in more detail, particularly regarding the branching from the metabolite oxaloacetate towards the reductive direction (MQO, FUM, and SDH) versus the oxidative direction (ACO and IDH) (**Fig. 6B**). The abundance of all these proteins in the reductive direction towards succinate and in the oxidative direction towards α-ketoglutarate were increased in the high*-n-*caprylate production condition (**Fig. 6B**). Interestingly, the abundances of the proteins (KOR and OOR) linking α-ketoglutarate to succinate were depleted in the high-caprylate treatment (**Fig. 6B**). The abundance of the protein (GLT) involved in the amination of α-ketoglutarate to produce glutamate in oxidative direction was positively correlated with *n*-caprylate production (**Fig. 6B**). The metaproteomics data, thus, implied that, rather than completing the carbon flow through the canonical oxidative side of the TCA cycle, the microbiome converted α-ketoglutarate into glutamate.

To investigate further the active metabolic routes in our bioreactor microbiomes, we performed a metabolomics profiling of the extra- and intracellular metabolomes for the long-term operated bioreactors.^38^ The metabolomics method surveyed for the detection of 250 detectable metabolites of which 25 were measurable and 11 were quantifiable (**Fig. 7A**). The 11 quantifiable metabolites were primarily TCA cycle intermediates (succinate, fumarate, and malate) or related to pathways directly connected to the TCA cycle (aspartate and pyroglutamate) (**Fig. 7A**). Amongst these quantifiable metabolites, succinate and pyroglutamate had 4-10-fold higher concentrations at one or both sampling times (**Fig. 7A**). Due to the higher *n*-caprylate output in both bioreactors on Day 937 than on Day 866, metabolites with a higher concentration on Day 937 were assumed to be possibly linked to *n*-caprylate production. Notably, the highest concentration was obtained for the extracellular pyroglutamate on Day 937 for both the AF and UASB reactors (**Fig. 7A**). Taken collectively the metaproteomics and metabolomics data, we propose a bifurcated TCA cycle with carbon fluxes in both directions in the reductive side (from oxaloacetate to succinate) and oxidative side (from oxaloacetate to α-ketoglutarate), thus explaining the accumulation of succinate and pyroglutamate derived from α-ketoglutarate (**Fig. 7B**).

**Fig. 7.**
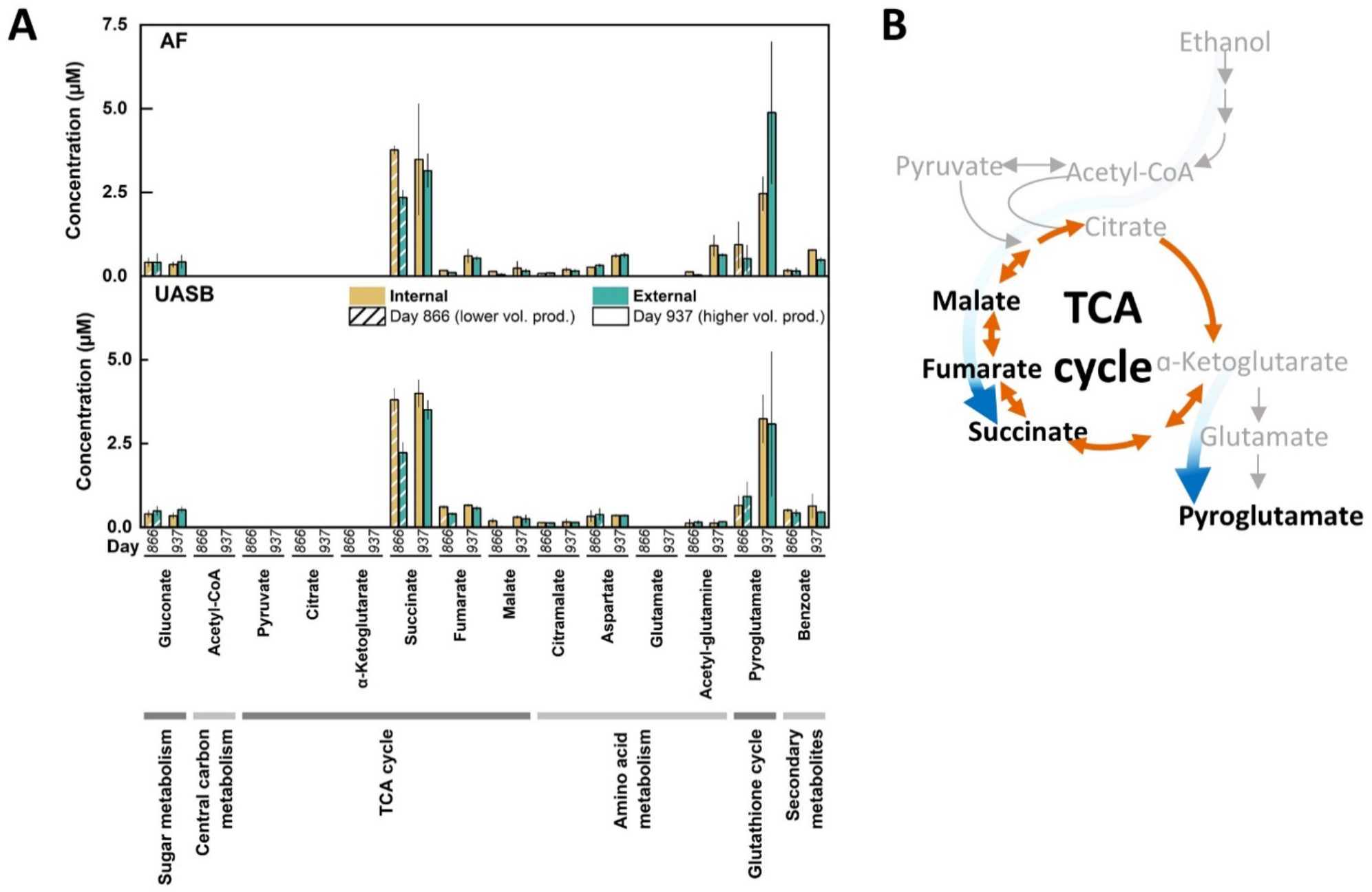
Measured metabolites and proposed metabolic routing from ethanol to intermediates based on metabolomics and metaproteomics data. **A**. The metabolite concentrations are shown as a grouped bar graph for the AF and the UASB reactors. From general to specific, the bars are grouped by metabolite, sampling time point (Day 866, 937), and intra-or extracellular concentration (Internal, External). The volumetric *n*-caprylate production rate of the reactors was higher on Day 937 compared to Day 866. We measured elevated concentrations of succinate and pyroglutamate. Succinate was found at higher concentrations in intracellular samples, regardless of the sampling timepoint. Pyroglutamate was found at higher concentrations on Day 937 than on Day 866 with no preference for intra-or extracellular samples. From the 250 metabolites included in the analysis, we acetyl-CoA, pyruvate, glutamate, phosphoenolpyruvate, α-ketogluconate, acetyl-glutamate, lactate, G6P, R5P, S7P, F6P, FBP, 6PG, 3PG at non-quantifiable concentrations. The error bars represent the standard deviation between technical triplicates. **B**. A schematic summary showing the proposed metabolic routing during the aerobic pre-conversion based on the metabolomics analysis. The blue arrows depict the flow of metabolites to either succinate or pyroglutamate. Quantified metabolites are highlighted in black and non-quantified metabolites in grey. Metabolite abbreviations: glucose-6-phosphate, G6P; ripose-5-phosphate, R5P; 3-phosphoglycerate, 3PG. sedoheptulose 7-phosphate, S7P; fructose-6-phosphate, F6P; fructose-1,6-bisphosphate, FBP; 6-phosphogluconate, 6PG; 3-phosphoglycerate, 3PG.

## Discussion

### Bioprocess operation to produce *n*-caprylate at high rates with an industrially applicable medium

During Period I, the AF reactor outperformed the UASB reactor in volumetric *n*-caprylate production rate and selectivity (**Fig. 1** and **Fig. S4B,D**, ESI †). The superior performance was likely due to improved retention of planktonic microbes and a more diverse community. We later noticed considerable biomass accumulation for the AF reactor upon removal of the packing material. Therefore, we concluded that the packing material acted as a cell retention system, decreasing cell washout. To the best of our knowledge, this study ranks third regarding the volumetric *n*-caprylate production rate at 0.14 g L^-1^ h^-1^ compared to the highest reported values of 0.33 g L^-1^ h^-1^ and 0.22 g L^-1^ h^-1^.^21, 22^ The three studies have an in-line product extraction system in common to remove toxic inhibiting *n*-caprylic acid efficiently.^21, 22^ In contrast to previous studies, we reached this value without ethanol and acetate accumulation in the fermentation broth (no overfeeding) and without adding yeast extract (we used a defined medium). The simplicity of the medium is crucial in reducing operating expenses when considering the scale-up of a bioprocess.^39^ The achieved *n*-caprylate selectivities were well above 50% for limited time intervals in both bioreactors, which is already high but needs further improvement to be commercially viable. This study identified oxygen supply as a critical factor, dictating volumetric *n*-caprylate production rates from ethanol and acetate.

### Ethanol is converted into intermediate metabolites in the presence of oxygen that enable *n*-caprylate production at high rates and selectivities

Through several different experiments, we discovered that oxygen depletion negatively affected the volumetric *n*-caprylate production rates: **(1)** when supplementing spikes of L-cysteine, which is an oxygen-scavenging molecule, into the medium during Period I, we observed a performance collapse in both bioreactors (**Fig. 1**); **(2)** the cumulative effect of oxygen-depleting modifications during Period II resulted in a process shutdown with an accumulation of unmetabolized substrate (**Fig. 1** and **Fig. S4A,C**, ESI †); and **(3)** in bottle experiments with stable-isotope tracing, the anaerobic treatment produced *n*-caprylate at sluggish rates and only half the final concentrations compared to the aerobic treatment (**Fig. 2C**).

Conversely, the presence of oxygen stimulated the production of *n*-caprylate. At first glance, this result contradicts the principle that chain elongation is a strictly anaerobic process.^19^ However, based on the results of the stable-isotope tracing bottle experiments, we hypothesized why the presence of oxygen would promote *n*-caprylate production. When examining the aerobic treatment, we observed carboxylate production after a delay of two days. However, cell densities increased drastically during the initial two days while ethanol concentrations decreased (**Fig. S5B,C**, ESI †). One of the reasons that the yield of carboxylates from consumed ethanol was lower for the aerobic treatment than the anaerobic treatment could be that microbes used the carbon for biomass growth when oxygen was present. High growth rates are typical for aerobic microbes, suggesting that they were involved in efficient ethanol uptake and oxygen consumption.^40^ When we set up the bottle experiments, we also administered a pool of intra- and extracellular metabolites from the fermentation broth inoculum. In fact, this is why the carboxylate yield for the anaerobic treatment was higher than 100% (*i.e*., 127%). We traced the introduced metabolites as part of the distributed fractional isotope abundances of carboxylates for the anaerobic treatment. The various fractional isotopomer abundances for the anaerobic treatment indicate that the ^13^C-labeled ethanol substrate and at least one introduced ^12^C-labeled metabolite were alternately incorporated (**Fig. 3B,D,F**).

If we again consider the aerobic treatment for the bottles, we cannot observe the same labeling patterns into carboxylates compared to the anaerobic treatment. Combining the evidence of the delayed onset of carboxylate production and the disappearance of labeling patterns, we identified that aerobic and/or facultative anaerobic microbes carried out a pre-conversion of ethanol into metabolites that are more suitable for the elongation from *n*-butyrate to *n*-caprylate than ethanol. The distributed labeling patterns are not visible in the aerobic treatment because labeled ethanol was not competing for integration with unlabeled metabolites (**Fig. 3A,C,E**). Instead, the aerobic treatment for the bottles contained ethanol and freshly aerobically formed metabolites, which were both labeled with heavy isotopes and integrated into carboxylates after oxygen depletion, which does not violate the principle of anaerobic chain elongation. Furthermore, the metabolites were favorably integrated into higher-chain carboxylates, evidenced by the higher total ^13^C-fraction of *n*-caproate and *n*-caprylate compared to *n*-butyrate for the aerobic treatment (**Fig. 2D-F**). This was the opposite for the anaerobic treatment for which the metabolite pool was unlabeled, explaining the reduced total labeling of the higher-chain carboxylates compared to *n*-butyrate by integration (**Fig. 2G-I**).

The sluggish rates of *n*-caprylate production for the anaerobic treatment in the bottles were due to the limited pool of these metabolites that were supplied with the fermentation broth as inoculum. The gradual increase of the ^13^C-fraction throughout the experimental period could be due to either suboptimal conditions for the metabolite synthesis for the anaerobic treatment or the slow integration of ethanol into *n*-caprylate.^29^ The ^13^C-fraction did not change substantially throughout the experimental period for the aerobic treatment, indicating that the ratio between ethanol and the metabolites was stable, and was not limited after the first two days of no carboxylate production. Indeed, ethanol was still present at the end of the 10-day experimental period for the aerobic treatment (**Fig. S5C**, ESI †). The rapid increase of *n*-caproate and *n*-caprylate for the aerobic treatment is a testimony to the potential of the observed metabolites for higher-chain carboxylate production. The plateauing of carboxylate concentrations after six days was likely due to the cytotoxicity of undissociated *n*-caproic acid and *n*-caprylic acid (**Fig. 2B-C**), emphasizing the need for product extraction.^23, 41^

Regarding the molecular assembly, we could predict possible chemical conversion routes based on the stable-isotope-tracing experiment. For the aerobic treatment in bottles, *n*-butyrate comprised slightly more than 75% (3/4 carbon atoms) ^13^C, which coincides with three carbon atoms per molecule (**Fig. 2D**). Also, the total ^13^C-fraction of *n*-caprylate was close to 87.5% (7/8 carbon atoms) (**Fig. 2F**). We, therefore, argue that *n*-butyrate was the starting compound for *n*-caprylate production after rapid ethanol conversion (one ^12^C per molecule). The elongation of the carbon chain from four to six progressed by the addition of two similarly labeled carbon atoms at a time at position 1. However, this current model did not hold for the elongation of *n*-caproate to *n*-caprylate after comparing the six-carbon-chain segments for these caboxylates. Instead, we observed a differently two-carbon-atom addition at position 7, which was not present in the anticipated location of positions 5 and 6 in *n*-caproate (**Fig. 4B**). Thus, the elongation of *n*-caproate to *n*-caprylate would follow a different, albeit unknown, mechanism. Our future work will ascertain this new metabolic pathway in chain-elongating microbiomes.

To support the findings of aerobically produced intermediate metabolites, we conducted two additional bottle experiments combining the knowledge gained during the stable-isotope tracing experiment, with the metabolomics profiling. We used thawed biomass, though, for these additional experiments. We measured potential intermediate metabolites for chain elongation that also occurred within the metabolomics profiling: succinate, lactate, pyroglutamate, and pyruvate. During the first additional bottle experiment, we compared again an aerobically and an an anaerobically inoculated treatment. We observed more rapid production of succinate and lactate paired with higher *n*-caproate specificity in the aerobic than in the anaerobic treatment (**Fig. S11**, ESI †).

In the second additional experiment, we compared two aerobic treatments, one with a reinoculation after 14 days, and one without reinoculation (**Fig. S12**, ESI †). The succinate concentration remained lower in the treatment with reinoculation than in the treatment without reinoculation while chain elongation to *n*-butyrate and *n*-caproate was higher. The exposure to oxygen in the first days likely reduced the ability of the microbiome to carry out anaerobic chain elongation after oxygen depletion.

The results of the additional bottle experiments support the hypothesis that oxygen intrusion: **(1)** promotes intermediate metabolite production to succinate and lactate which in turn accelerate chain elongation; **(2)** the anaerobic community needs to be strong and protected from oxygen intrusion, as observed during the second additional experiment were the reinoculation showed higher *n*-caproate production. During the additional bottle experiments only low concentrations of *n*-caprylate (∼ 0.8 mM C) were produced within the aerobic treatment of the first reiteration. The reduced *n*-caprylate production and overall slow chain elongation was most likely caused by the thawed inoculum that was not as vital as the fresh inoculum, that we had used for the stable-isotope tracing bottle experiments.

### A trophic hierarchy of aerobic pre-conversion to stepwise anaerobic chain elongation enabled the production of *n*-caprylate at high rates

We postulate a trophic hierarchy within the reactor microbiomes by combining the chemical evidence of the metabolite formation for the aerobic treatment in bottles with the data from metagenomics, metaproteomics, metabolomics, and the isolation of a physiologically novel *C. kluyveri* strain (**Fig. 8**). The core-community structure suggests that strictly aerobic bacteria, which were represented predominantly by *P. caeni*, or facultative anaerobic fungi, carried out a pre-conversion of ethanol into intermediate metabolites (**Fig. 5**). Previously, *P. caeni* was present in several chain elongation studies using alcohols as feedstock. Some studies identified *P. caeni* according to its family Micrococcaceae or order Micrococcales.^21, 22, 42-48^

**Fig. 8.**
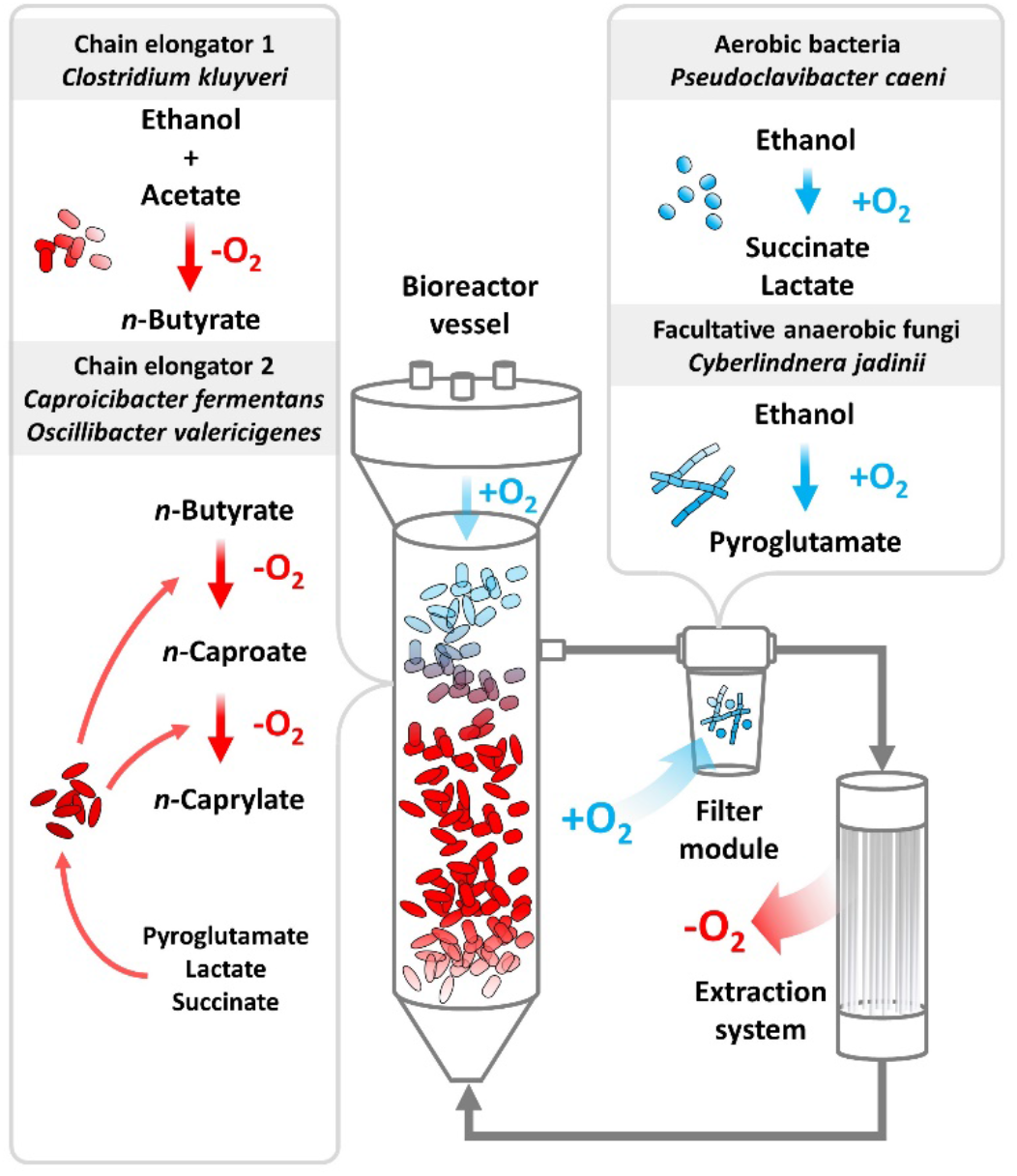
Schematic depiction of oxygen intrusion into the bioreactors during Period I. The liquid flow is shown from the medium tank through the bioreactor vessel to the fermentation broth recirculation. The oxygen intrusion into the filter module and headspace of the bioreactor vessel is illustrated as blue arrows. The oxygen removal in the extraction system is illustrated as red arrow. The filter module was a source of oxygen, and the extraction system was a sink of oxygen. The proposed main reactions from the substrate ethanol and acetate to the desired product *n*-caprylate are schematized. Aerobic bacteria (*e.g., P. caeni*) or facultative anaerobic fungi (*e.g., C. jadinii*) converted a part of the ethanol into succinate, lactate, and pyroglutamate. Subsequently, the first chain elongator (*C. kluyveri*) elongated acetate with the residual ethanol into *n*-butyrate. Next, the second chain elongators (*C. fermentans* and *O. valericigenes*) elongated *n*-butyrate with succinate, lactate, or pyroglutamate to *n*-caproate and then *n*-caprylate. The reactions in the presence of oxygen occurred in the filter module and partly in the headspace of the bioreactor vessels. During Period II, we removed the filter module and sparged the headspace with nitrogen, stopping aerobic pre conversion.

Combining the metabolomics and metagenomics data, we wondered about the chemical nature of the metabolites and the microbial species producing it. The metabolite profiling showed succinate and pyroglutamate accumulation in the fermentation broth and our additional bottle experiments verified lactate production (**Fig. 7A** and **Fig. S11, S12**, ESI †). Previously, a study associated succinate formation in chain elongating microbiomes with the phylum Actinobacteria.^49^ The most abundant aerobic bacterium in our food web, *P. caeni*, belongs to the Actinobacteria phylum, which produces succinate and lactate. Another member of our core community, the facultative anaerobic fungus *C. jadinii*, produces pyroglutamate under conditions of oxidative stress (**Fig. 5, Fig. 8**).

While we cannot pinpoint, which metabolite was responsible for increased chain elongation with oxygen intrusion, the metaproteomics data and metabolomics profiling suggest that pyroglutamate is an important intermediate metabolite. Within the metaproteomics data we found the phosphoserine-aminotransferase, which utilizes glutamate, of *O. valericigenes* only in the high *n*-caprylate production-rate treatment. Additionally, we noticed an increase in pH during the assumed metabolite conversion during the first two days for the aerobic treatment in bottles (**Fig. S5A**, ESI †). Ammonia is a side product of glutamate conversion, which acts alkaline and can be a plausible reason for the increase in pH.^50^

Previous studies that employed fully anaerobic microbiomes mentioned reduced carboxylate conversion efficiencies with ethanol concentrations exceeding 10 g L^-1^. Here, the ethanol concentration in the medium was 13.42 g L^-1^. We, therefore, conclude that aerobic pre-conversion of ethanol into less toxic metabolites was essential to maintain high volumetric production rates for *n*-caprylate.

While a certain amount of oxygen was crucial to the process performance, the correct dosing proved to be a sensible parameter. During Period IV, we first sparged air at a comparatively low flow rate to the bottom of the bioreactor vessels. The residual ethanol concentration in the fermentation broth decreased due to the efficient ethanol conversion of strictly aerobic and/or facultative anaerobic microbes. Nevertheless, chain elongation to higher-chain carboxylates was transient and sluggish (**Fig. 1**). The product spectrum shifted from mainly *n*-caprylate during Period I to mostly *n*-butyrate with some *n*-caproate and *n*-caprylate during Period IV, after air sparging (**Fig. 1**). We conclude that the strict aerobic bacteria and facultative anaerobic fungi did not remove enough oxygen from the fermentation broth to allow high-rate anaerobic chain elongation to *n*-caprylate. Toward the end of the operating period, we increased the airflow into the bioreactor vessels, which let anaerobic chain elongation fully collapse (L in **Fig. 1**). As implied by the transient volumetric production rates during Period IV, aeration of the bioreactor vessels created a mixed environment in which aerobic ethanol uptake and anaerobic chain elongation proceeded side by side. Spatial segregation of aerobic and anaerobic conversions led to more optimal process performances, as observed during Period I. During this period, much of the oxygen intrusion occurred only through the top layer of the liquid column *via* the headspace or through the filter modules. These filter modules contained a distinct microbial species arrangement, which were enriched in aerobic bacteria and facultative anaerobic fungi (**Fig. 5**, and **Fig. 8**). Still, our bioreactor setup leaves much room for improvement regarding oxygen supplementation,^51^ because we did not optimize or design it with oxygen intrusion in mind. The production rates of *n*-caprylate fluctuated the most from the produced carboxylates, caused by suboptimal management of aerobic and anaerobic phases. Additionally, we noticed low biomass concentration in the bioreactors, which was evident by a relatively clear fermentation broth on top of a small amount of anaerobic granules in the UASB reactor. A resulting high volumetric ratio of fermentation broth to biomass could indicate that: **(1)** the observed dilution is preferable due to the cytotoxic effects of metabolites or products; or **(2)** the process parameters could be optimized, and we could increase the volumetric production rate by selecting more suitable operating conditions.

During operating periods when *n*-caprylate production in the bioreactors faded, we observed a surge in ethanol and *n*-butyrate concentration in the fermentation broth (*e.g*., starting Day 645) (**Fig. 1, Fig. S4A,B**, ESI †). The ethanol concentration increased because the aerobic metabolite production stopped. Without the metabolites, *n*-caprylate production stagnated. As a result, *n*-butyrate was not integrated into *n*-caprylate and accumulated in the fermentation broth. This causative sequence also explains the negative correlation of *C. kluyveri* to *n-*caprylate production. More ethanol was available when the aerobic pre-conversion to the intermediate metabolites stopped. Ethanol is the electron donor for *C. kluyveri*, from which it produces *n*-butyrate. Therefore, the relative abundance of *C. kluyveri* increased parallel to rising ethanol concentrations until the ethanol reached toxic concentrations. In our hypothetical food web, *C. kluyveri* was the first trophic group that performed chain elongation (**Fig. 8**). For two additional chain-elongation steps, the second trophic group of chain elongators, *C. fermentans*, and *O. valericigenes*, converted *n*-butyrate to *n*-caprylate (**Fig. 5, Fig. 8**).

The two-carbon-atom additions of chain elongation suggest acetyl-CoA-based metabolic pathways. Indeed, reverse ß-oxidation, which is known as a two-carbon-atom addition, was the predominant metabolic pathway in our reactor microbiomes (**Fig. 6A**). Nevertheless, other carbon chain-altering pathways of carboxylates, such as the TCA cycle or fatty acid degradation, may also play a role in the assembly of *n*-caprylate. The high relative abundances of hydrogenotrophic methanogenic archaea imply that hydrogen generation plays a substantial role in the metabolic pathways. In fact, reducing the hydrogen partial pressure by these methanogens likely increased *n*-caprylate production rates by lowering product inhibition (hydrogen is a side product of reverse ß-oxidation). In addition, this also considerably reduces the pertinence of the fatty acid biosynthesis pathway as a carbon chain-altering pathway of carboxylates because it has no hydrogen-producing step. The fatty acid biosynthesis pathway is an anabolic process and does not produce the required ATP. Finally, our metaproteomics analysis showed a very low total abundance of the proteins for the fatty acid biosynthesis pathway and a downregulation during high volumetric *n*-caprylate production rates (**Fig. 6A**). Therefore, our study concludes that the fatty acid biosynthesis pathway is not contributing to chain elongation. Furthermore, the chain elongation rates of our bioreactors were not limited by excessive ethanol oxidation, as described in previous studies.^18^ On the contrary, when employing high ethanol concentrations, its conversion into more potent intermediate metabolites is desirable to lower toxicity and increase production rates.

## Conclusion

We identified oxygen supplementation as a critical parameter for *n*-caprylate production with bioreactor microbiomes from high concentrations of ethanol and acetate *via* chain elongation. We found a trophic hierarchy that explains the oxygen dependency of the process. Bottle experiments showed that aerobically produced metabolites enable high-rate *n*-caprylate production. Based on extensive metabolite profiling, we hypothesize that the observed metabolite is succinate, lactate, and/or pyroglutamate, or a mixture of those. *P. caeni* and the facultative anaerobic fungus *C. jadinii* initiate the trophic hierarchy, converting ethanol into the intermediate metabolites in the presence of oxygen. *C. kluyveri* produced solely *n*-butyrate under anaerobic conditions, which we proved by isolating a physiologically novel strain from our bioreactor. The anaerobic bacteria *C. fermentans* and *O. valericigenes* used succinate, lactate, or pyroglutamate to elongate *n*-butyrate to *n*-caprylate with *n*-caproate as an intermediate. Stable-isotope-tracing experiments revealed that the recent integration step into *n*-butyrate and *n*-caproate occurred with two-carbon-atom integrations at position 1, suggesting acetyl-CoA-based pathways. However, the labeling pattern of *n*-caprylate was different. It showed a disintegration of the similarly-labeled two-carbon-atom integration at the carbon atoms in positions 7 and 8, suggesting differently labeled integration at the other side of the carboxylate, contrary to what the current model predicted. However, we do not know the mechanism. We used the metaproteomics data to deduce transparent relationships between metabolic pathways and *n*-caprylate production rates. Overall, reverse ß-oxidation was the most abundant and the TCA cycle the most positively correlated metabolic pathway to *n*-caprylate production. The fatty acid biosynthesis pathway was not related to chain elongation. With the knowledge of this microbial food web for *n*-caprylate production, we are now developing an industrial production platform based on chain elongation.

## Supporting information

Manuscript

## Author Contributions

All authors have approved the final version of the manuscript.

**Conceptualization**: K.G., B.S.J., L.T.A.; **Data Curation**: K.G., B.S.J., J.N.N; **Formal Analysis**: K.G., B.S.J., J.N.N, T.N.L., N.N., N.Z.; **Funding Acquisition**: L.T.A.; **Investigation**: K.G., B.S.J., J.N.N, H.W., C.S., I.B., N.N., N.Z.; **Methodology**: K.G., B.S.J., J.N.N, T.N.L.: I.B., N.N., N.Z., J.G.U., L.A.; **Project Administration**: K.G., B.S.J., J.N.N, L.T.A.; **Resources**: D.H.H., R.B.H.W., B.M., L.A., L.T.A.; **Software**: K.G., T.N.L., N.N; **Supervision**: J.G.U., D.H.H., R.B.H.W., L.A., L.T.A.; **Visualization**: K.G.; **Writing – Original Draft**: K.G.; **Writing – Review & Editing**: K.G., B.S.J., J.N.N, H.W.,C.S., T.N.L., N.N., R.B.H.W., L.A., L.T.A..

## Data Availability

The data supporting this article have been included as part of the Supplementary Information.

## Acknowledgements

**L.T.A**. acknowledges the support from the Alexander von Humboldt Foundation in the Alexander von Humboldt Professorship framework endowed by the Federal Ministry of Education and Research in Germany and the Novo Nordisk Foundation CO_2_ Research Center with grant number NNF21SA0072700. This work was supported by the Cluster of Excellence “Controlling Microbes to Fight Infections” (CMFI) and the German Research Foundation (DFG: EXC 2124 – 390838134) to **L.T.A** and **D.H**.. **H.W**. acknowledges the support from the China Scholarship Council (CSC). **I.B**. and **R.B.H.W**. acknowledge the support from the Singapore National Research Foundation and Ministry of Education under the Research Centre of Excellence Programme. Funding for **N.Z**. was provided by the U.S. Department of Energy (DE-SC0022181). **L.A**. acknowledges a Bessel Research Award from the Alexander von Humboldt foundation in support of her sabbatical time in Germany. Metabolomics analysis was funded by a CAREER grant awarded to **L.A**. from the U.S. NSF (NSF-CBET-1653092). We thank Raphael Mayer from the University of Tübingen for his support in operating the bioreactors.

