## Supplementary material for "Toward industrial C8 production: Oxygen intrusion drives renewable *n*-caprylate production from ethanol and acetate *via* intermediate metabolite production": Manuscript

Electronic Supplementary Information

### Supplementary Figures

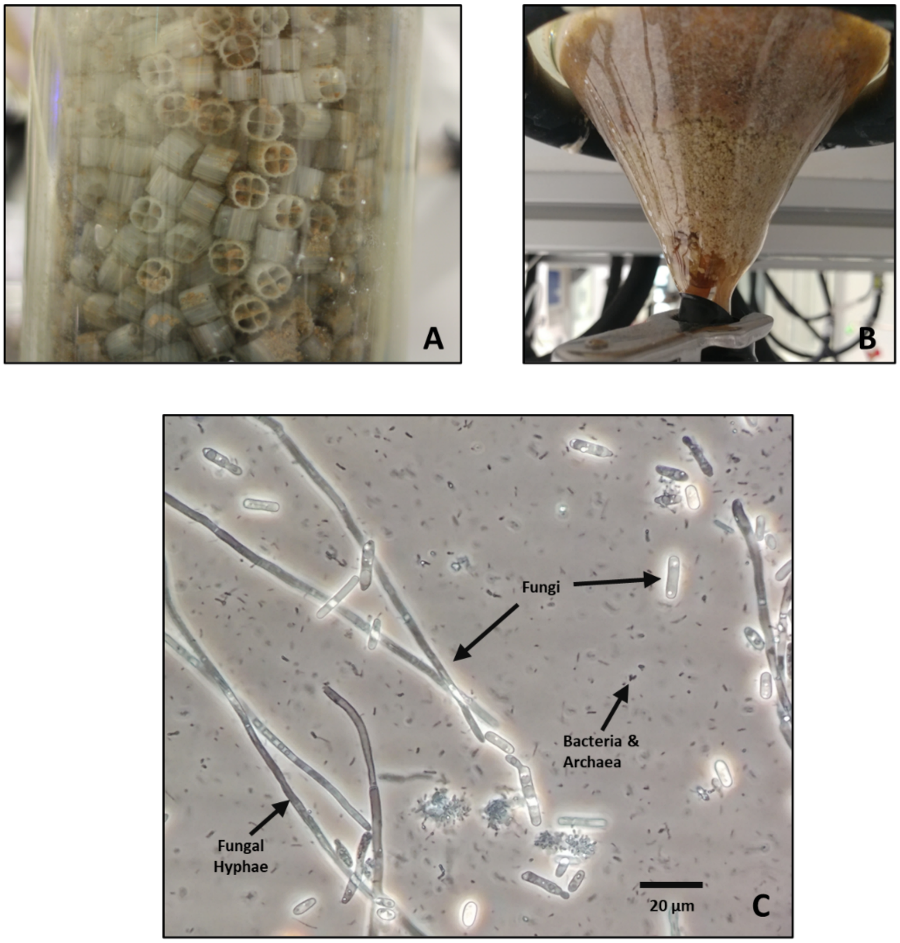


**Fig. S1.** Explanatory views of biomass accumulation **A.** inside the anaerobic filter (AF) reactor and **B.** in the bottom of the upflow anaerobic sludge blanket (UASB) reactor. In the AF reactor, biomass accumulated mainly on the surface of the wheel-shaped packing material by biomass trapping and settling. In the UASB reactor, biomass accumulated as granules at the bottom of the reactor vessels. The microbial community **C.** on Day 419 of the operating period under 40 x magnification depicts large fungi with hyphae. The smaller black cells are bacteria and methanogenic archaea.

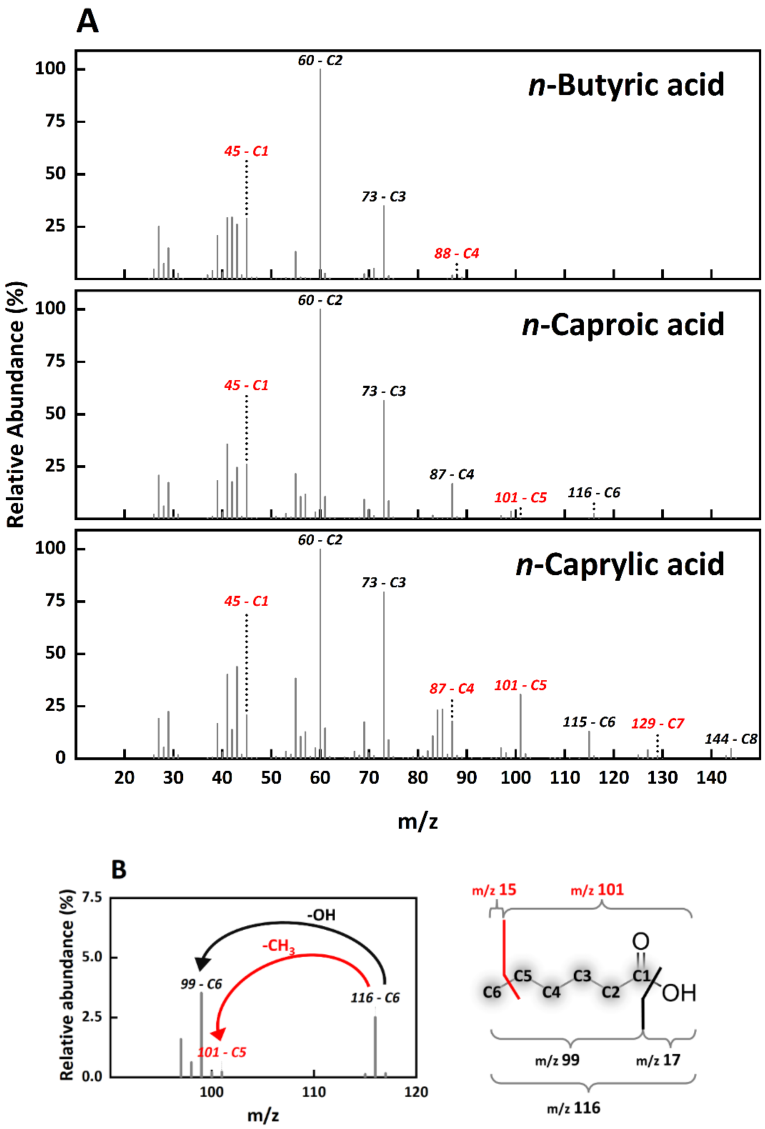


**Fig. S2.** Explanatory figure of mass spectra that we obtained with gas chromatography coupled to mass spectrometry. The carbon-chain segments from positions 1-8 are marked as C1-C8 with the respective atomic weight (*m/z* – position). The direction of view starts from the functional group, and the segment includes all lower-numbered carbon atoms up to the marked number. **A.** The mass spectra of measured unlabeled *n*-butyrate, *n*-caproate, and *n*-caprylate standards are shown. The utilized mass spectrometer produces mass spectra by electron ionization, which is a hard fragmentation method. In the ionization source of the mass spectrometer, electrons are accelerated to 70 eV and collide with the analyte. As a result, the analyte fragmented into multiple positively charged ions that were detected. The fragmentation pattern is indicative of the analyte and depends on the stability of the chemical bonds of the investigated chemical and the fragmented ions. Tags indicate the mass and carbon-chain segment in the mass spectra.The base peak is in all cases at *m/z* 60 and set to 100% relative abundance.[1] The remaining peaks are adjusted to a relative abundance proportional to the base peak. The parent peak of the whole molecular ion is visible in all three cases. Black tags indicate carbon-chain segments from which the labeling pattern can be derived. Red tags show carbon-chain segments from which the labeling pattern cannot be derived due to overlap with the labeling pattern of other fragments. **B.** An example of overlap is shown with the zoomed-in mass spectrum of *n*-caproate on the left side. The peak at *m/z* 116 resembles the whole *n*-caproate carbon-chain segment, which is illustrated on the right side. The peak at *m/z* 99 is the acylium ion (caproyl-ion), which is formed by cleaving the hydroxy group from *n*-caproate. The peak at *m/z* 99 is considerably higher than the peak at *m/z* 101, which resembles the cation derived from cleaving the carbon in position 6. By labeling the peak at *m/z* 99 more than once with $^{13}$C-carbon, the relative abundance of the acylium ion overlaps the five-carbon-chain segment peak at *m/z* 101. Therefore, the labeling pattern of the five-carbon-chain segment of *n*-caproate cannot be inferred.

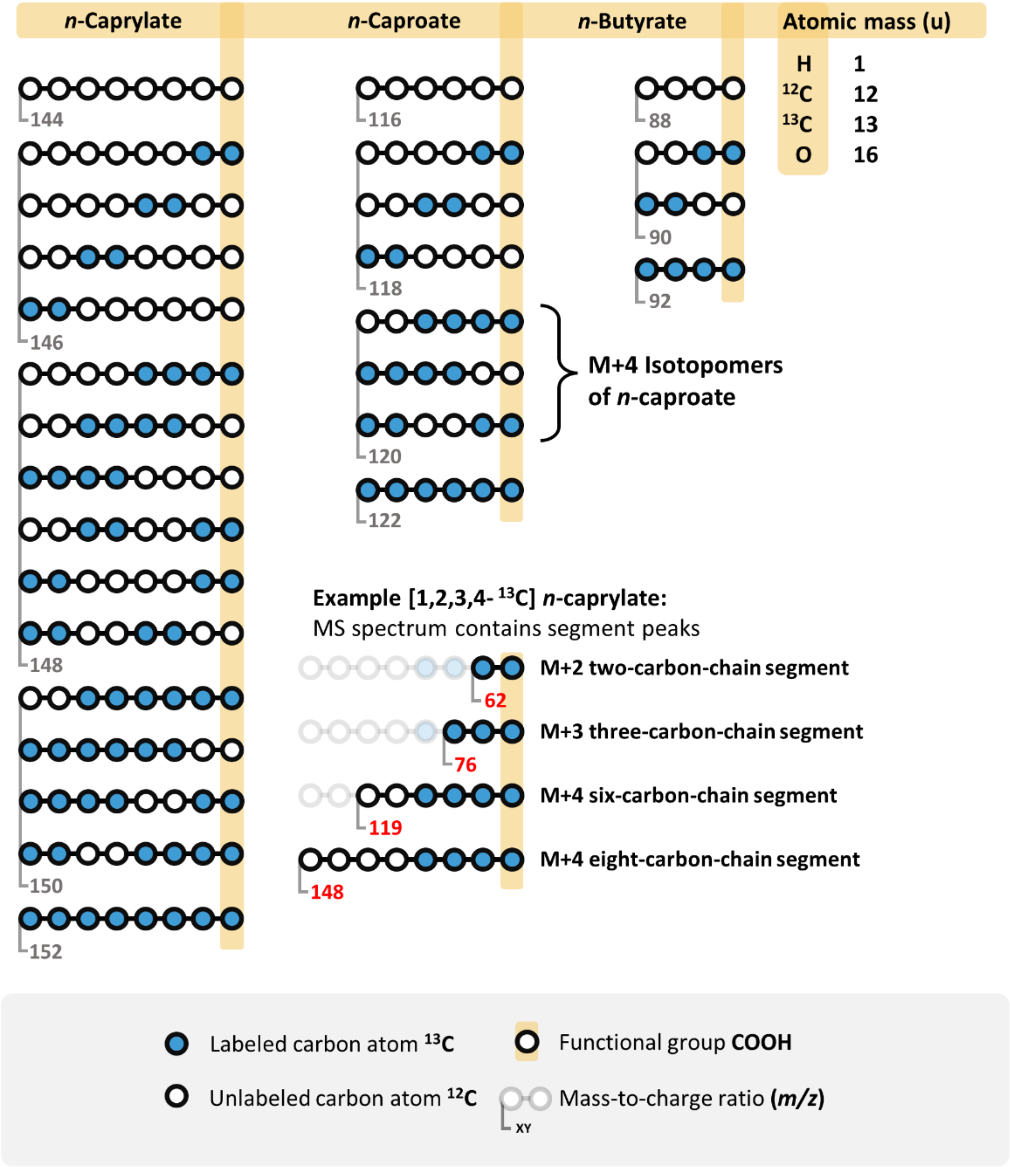


**Fig. S3.** Labeling examples, assuming two-carbon-step addition into carboxylates at the same time. Isotopomers are molecules with the same molecular formula and isotopic composition (same mass) but different spatial arrangements of these isotopes. In this case, the spatial arrangement of $^{13}C$ and $^{12}C$ differs. Isotopomers of each carboxylate are grouped. Using the described method, the example of [1,2,3,4-$^{13}C$] *n*-caprylate shows the obtainable carbon-chain segments. Notably, the entire molecule of *n*-caproate and the six-carbon chain of *n*-caprylate have an *m/z* value difference of one at the same labeling extent. The difference emerges because the six-carbon chain of *n*-caprylate is cleaved from the carbon atoms in positions seven and eight. As a result of the cleavage, the third hydrogen atom in position six is missing, leading to an *m/z* value of one less.

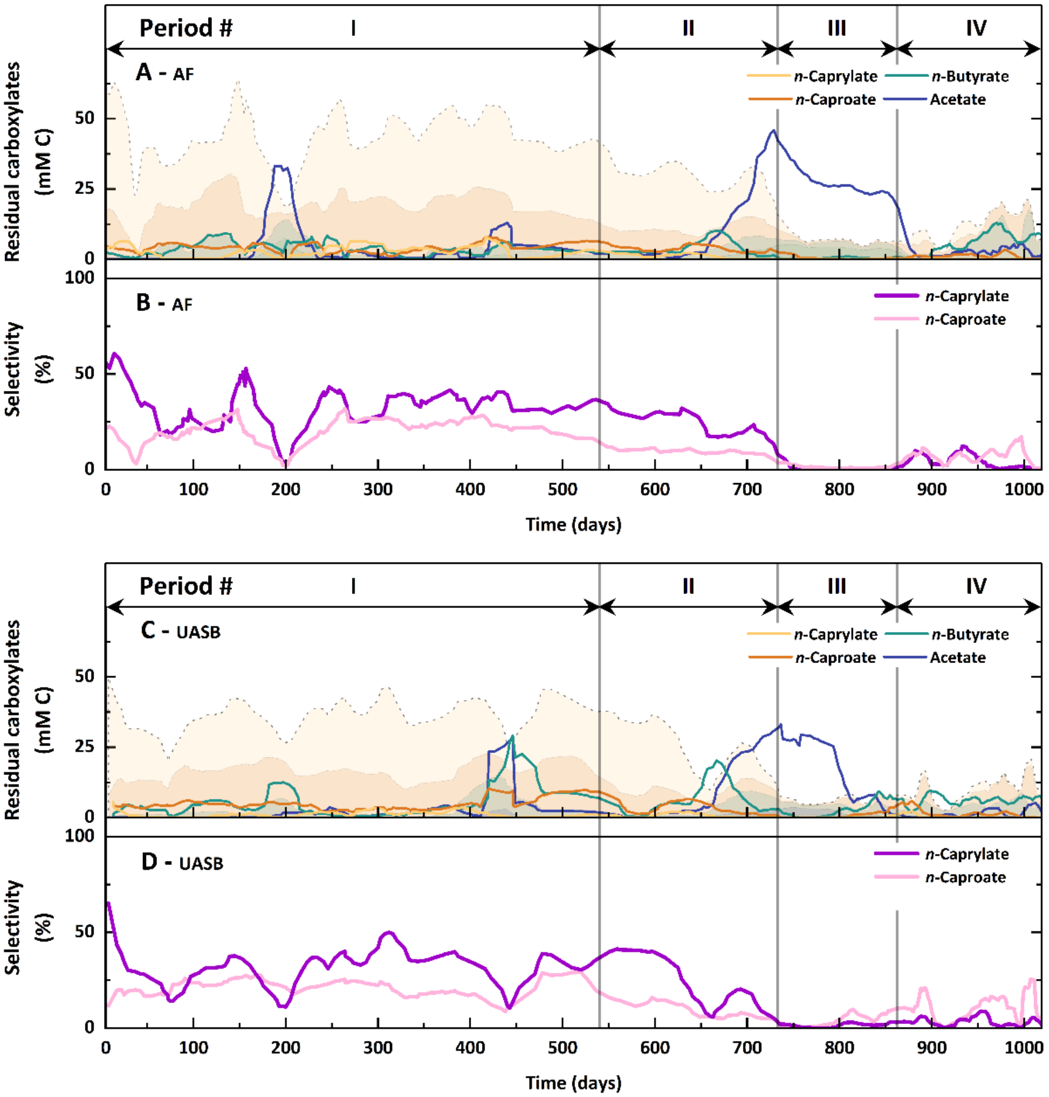


**Fig. S4.** Residual carboxylates in the fermentation broth and selectivity (% mol C/mol C, specific product/total substrate) into *n*-caproate and *n*-caprylate of **A, B**. the AF reactor and **C, D**. the UASB reactor throughout the operating period. The residual carboxylates (mM C) and the selectivity (%) were calculated as a six-measurement moving average. In Period I, when L-cysteine was added (Day 174-194 and Day 419-449), both bioreactors showed sharp drops in carbon selectivities. Additionally, during Period III, the efficiencies were close to zero when all oxygen supply was cut off. Lower efficiencies of *n*-caprylate production corresponded to an accumulation of either acetate or *n*-butyrate in the fermentation broth. Values were determined from samples collected daily or every other day. **A,C.** The faint area in the back of the residual carboxylates figure illustrates the bioreactor performance.

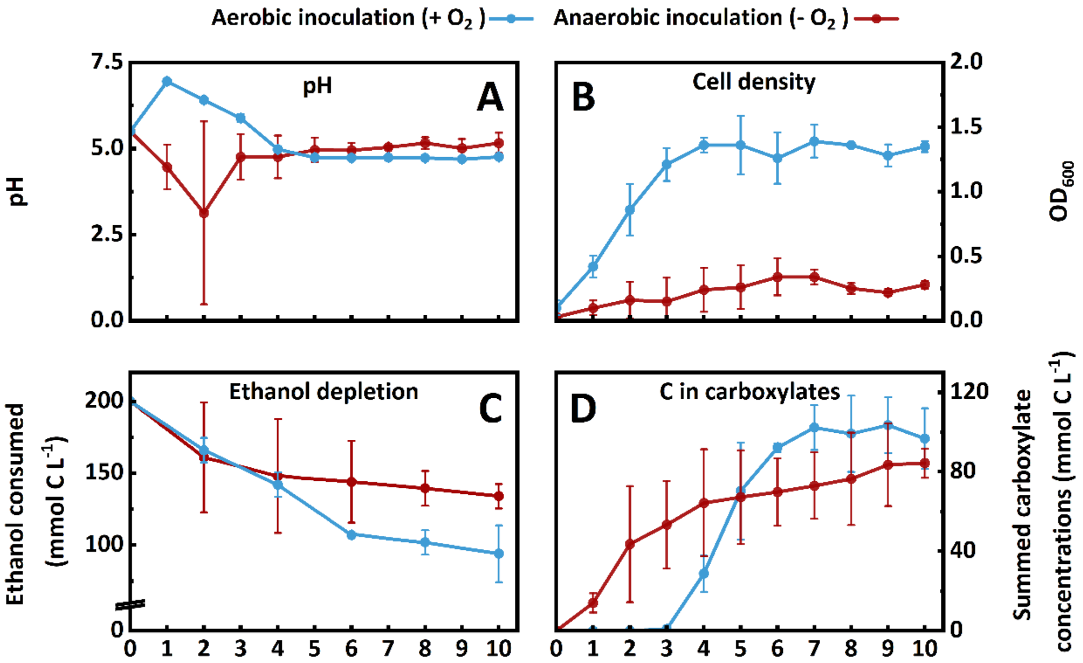


**Fig. S5. A.** pH, **B.** cell density, **C.** ethanol depletion, and **D.** total carbon (C) in carboxylates throughout the experimental period of 10 days for the batch experiments. The anaerobically inoculated bottles are depicted in red, and the aerobically inoculated bottles are light blue. **A.** The pH settled around 5.0, close to the pKa (~4.8-4.9) of produced carboxylic acids after a diverging beginning phase. **B, C.** The cell density ($OD_{600}$) and the ethanol depletion (mM C) show higher growth and more ethanol depletion in the aerobically inoculated bottles. **D.** The total carbon in carboxylates (mM C) shows production from Day 1 in the anaerobically inoculated bottles and a slow increase. The aerobically inoculated bottles show a delayed production onset with rapid growth and plateauing. The depicted values derive from daily sampling, and the errors represent the standard error between biological duplicates.

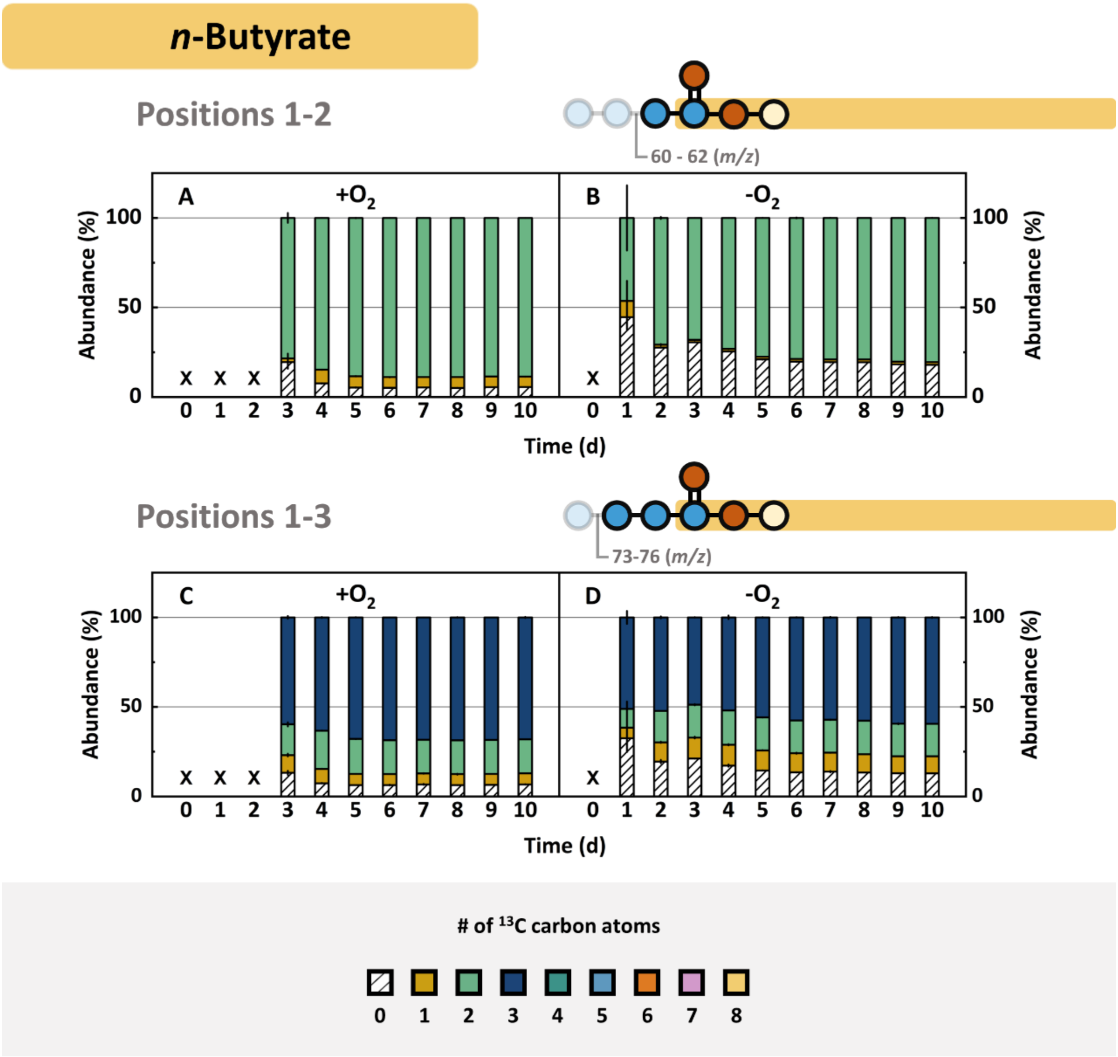


**Fig. S6**. Fractional isotopomer abundances of the **A,B.** two and **C,D.** three-carbon-chain segments of *n*-butyrate. The one- and four-carbon-chain segments were not analyzable due to overlapping signals with other fragments (*n*-butyric acid in **Fig. S2A**). Each graph shows fractional isotopomer abundances of the aerobic inoculated bottles on the left (+$O_2$). The data regarding anaerobically inoculated bottles is on the right (-$O_2$). The isotopes are arranged from bottom to top by increasing $^{13}C$ content. The depicted values derive from daily sampling, and the errors represent the standard error between biological duplicates.

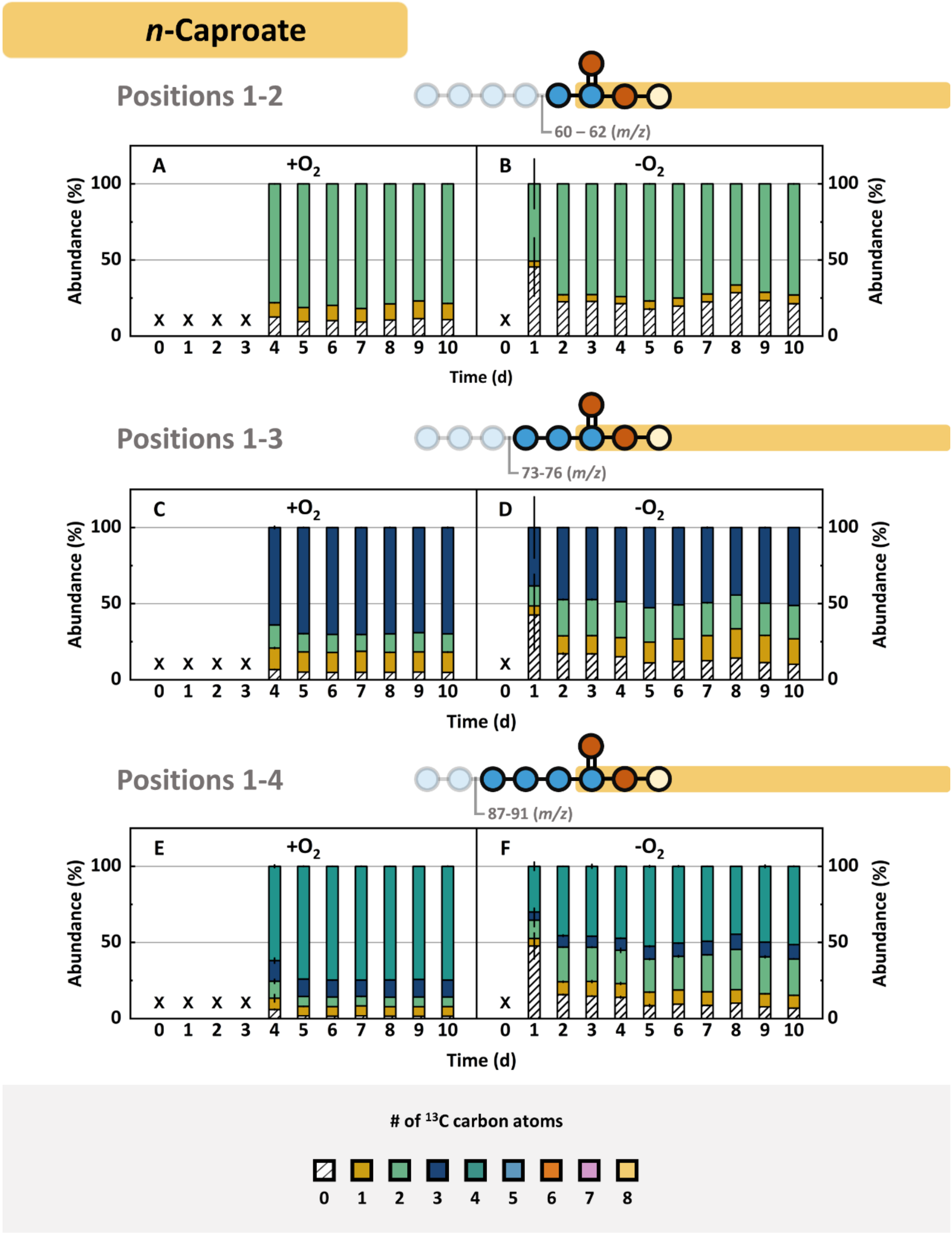


**Fig. S7** Fractional isotopomer abundances of the **A,B.** two-, **C,D.** three-, and **E,F.** four-carbon-chain segments of *n*-caproate. The one and five-carbon-chain segments were not analyzable due to overlapping signals with other fragments (*n*-caproic acid in **Fig. S2A**). Each graph shows fractional isotopomer abundances of the aerobic inoculated bottles on the left (+$O_2$). The data regarding anaerobically inoculated bottles is on the right (-$O_2$). The isotopes are arranged from bottom to top by increasing $^{13}C$ content. The depicted values derive from daily sampling, and the errors represent the standard error between biological duplicates.

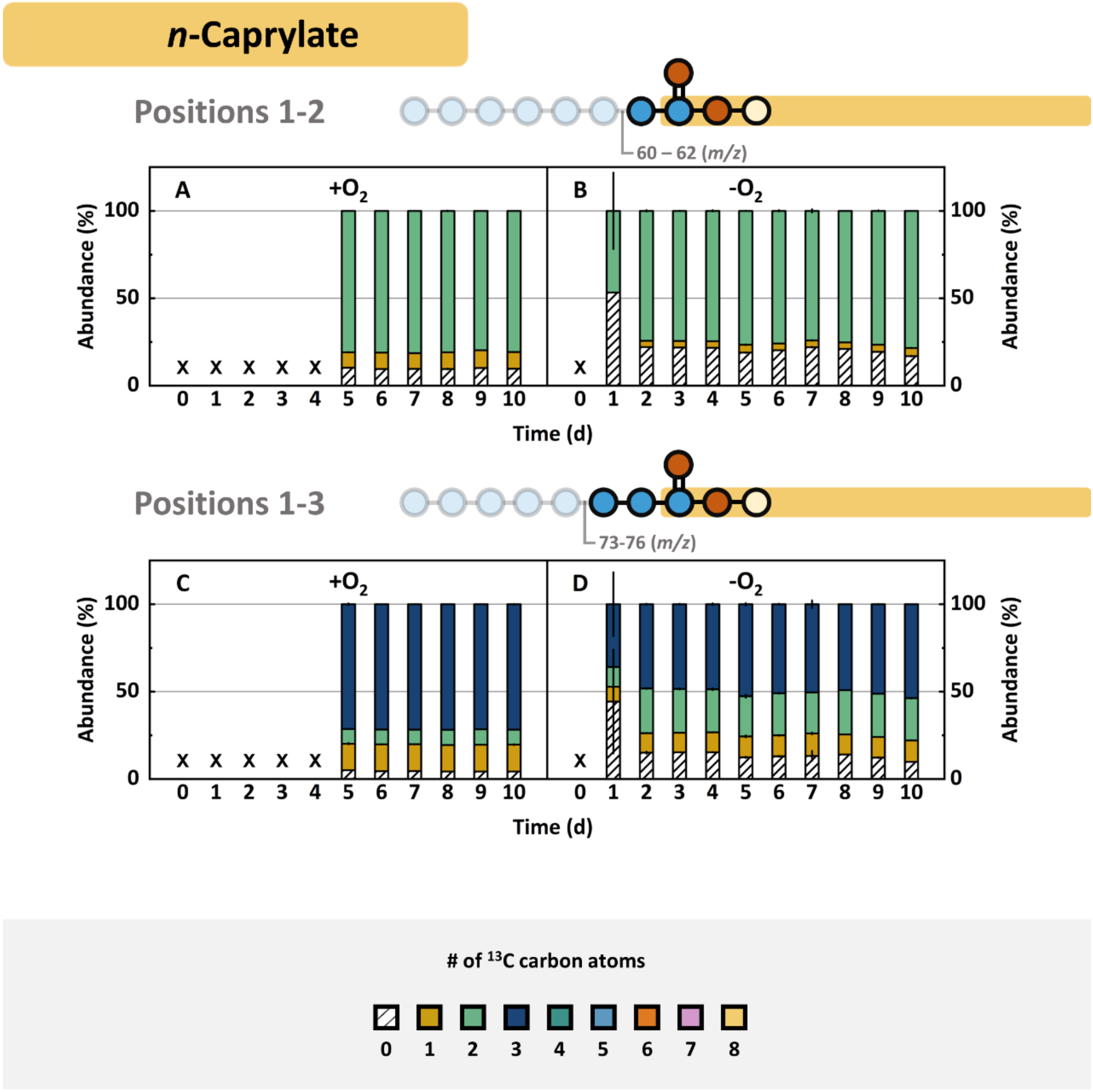


**Fig. S8** Fractional isotopomer abundances of the **A,B.** two- and **C,D.** three-carbon-chain segments of *n*-caprylate. The one, four-, five-, and seven-carbon-chain segments were not analyzable due to overlapping signals with other fragments (*n*-caprylic acid in **Fig. S2A**). Each graph shows fractional isotopomer abundances of the aerobic inoculated bottles on the left (+$O_2$). The data regarding anaerobically inoculated bottles is on the right (-$O_2$). The isotopes are arranged from bottom to top by increasing $^{13}C$ content. The depicted values derive from daily sampling, and the errors represent the standard error between biological duplicates.

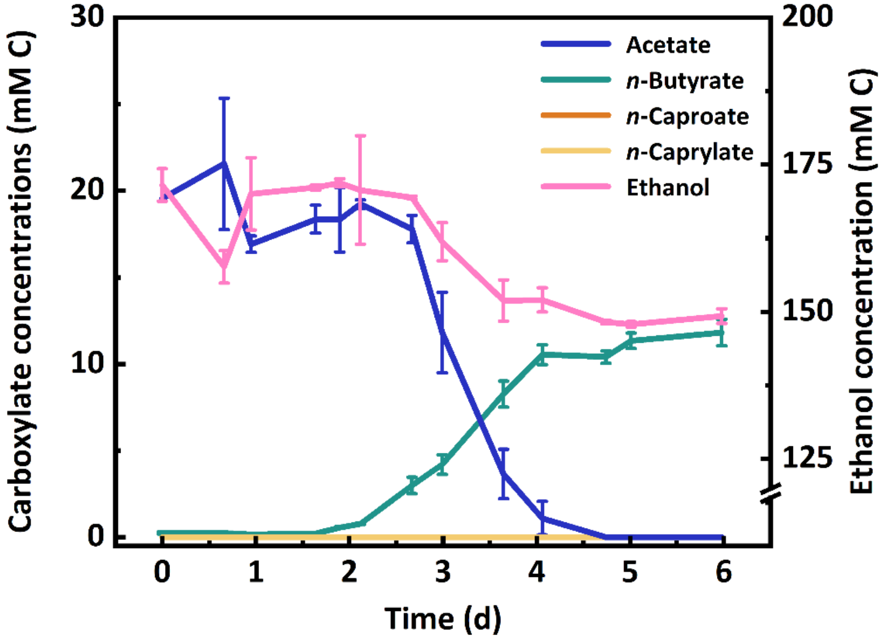


**Fig. S9.** Carboxylate and ethanol concentrations throughout six days lasting batch experiments with *Clostridium kluyveri* isolated from the AF reactor. The isolate produced exclusively *n*-butyrate among the carboxylates from the second day. The concentrations of *n*-caproate and *n*-caprylate did not exceed zero. Acetate and ethanol were constituents of the medium and readily depleted. The batch experiments were performed in biological triplicates. The error bars represent the standard deviation between the biological triplicates.

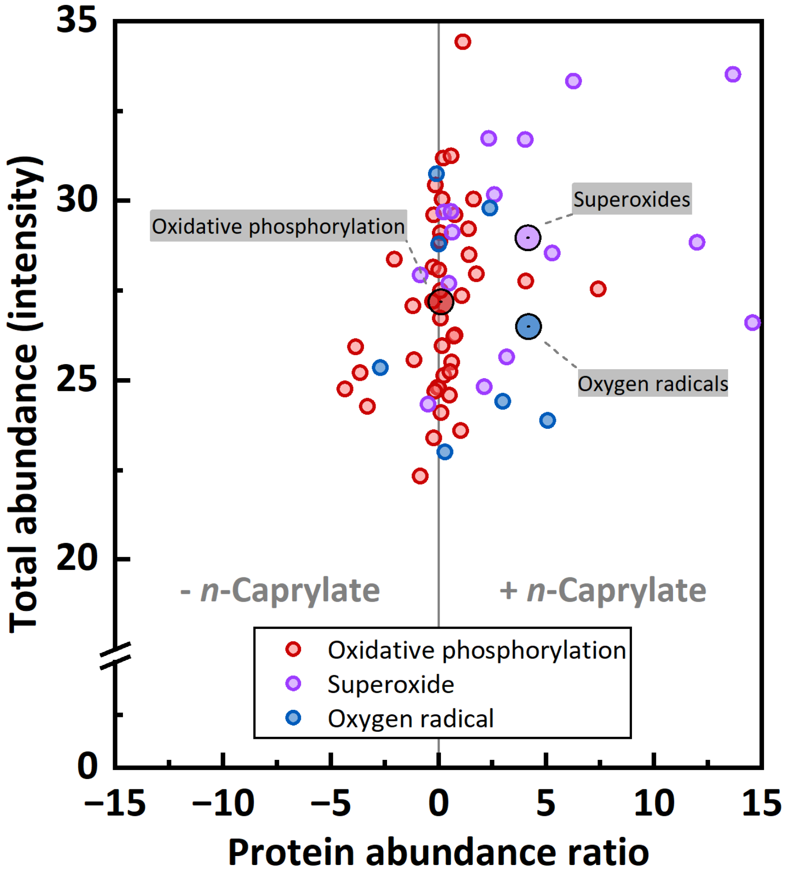


**Fig. S10.** Total abundances and the ratio of proteins related to oxygen presence. Proteins were mapped on the oxidative phosphorylation pathway and their function as oxygen-detoxifying agents. The high volumetric *n*-caprylate production-rate phase (+ *n*-Caprylate) was during regular operation, and the low volumetric *n*-caprylate production-rate phase (- *n*-Caprylate) was achieved by adding L-cysteine to the bioreactor medium (treatment). Oxidative phosphorylation was weakly positively related to the volumetric *n*-caprylate production rate. Other terminal electron acceptors, such as sulfate or nitrate, may have been abundant. Proteins related to treating superoxides and oxygen radicals were positively related to the volumtric *n*-caprylate production rate, confirming that oxygen intrusion was indeed related to the high *n*-caprylate production rate. The big circles represent average values for protein groups.

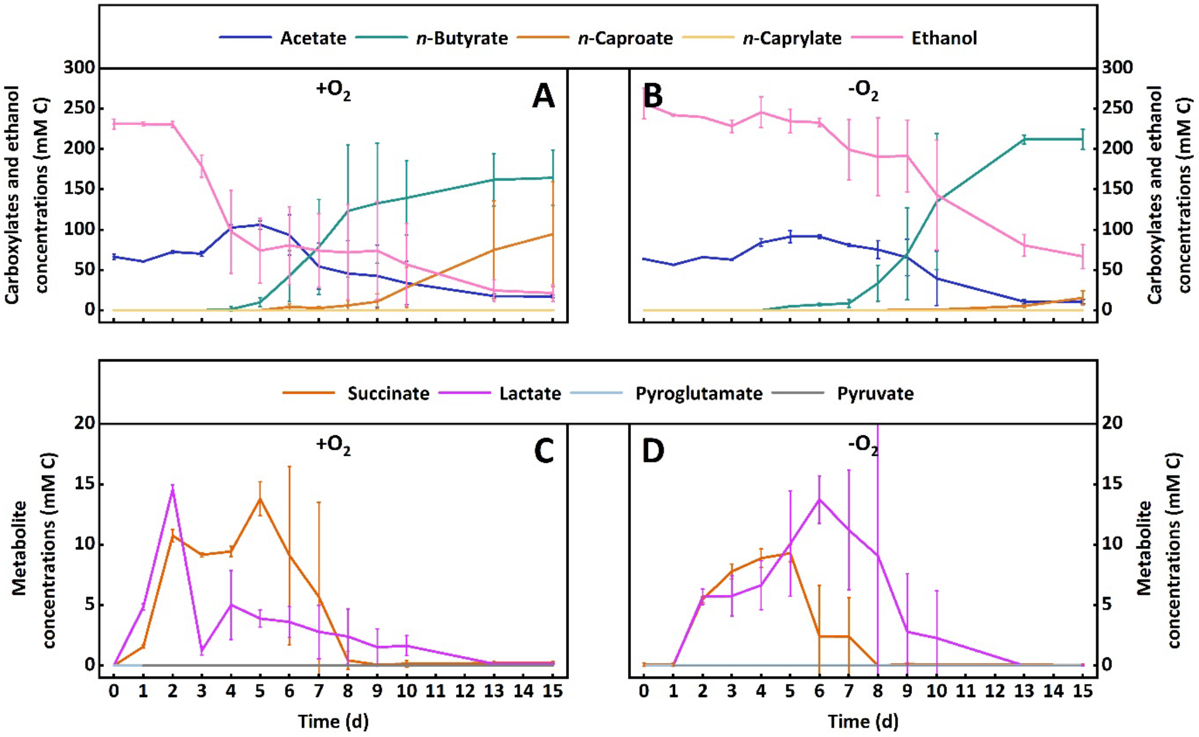


**Fig. S11.** Substrate, carboxylate (**A.** and **B.**), and intermediate metabolite (succinate, lactate, pyroglutamate, and pyruvate) (**C.** and **D.**) concentrations for the first additional bottle experiment throughout the 15-day experimental period. The aerobic treatment is shown on the left figures (+$O_2$) and the anaerobic treatment on the right figures (-$O_2$). The ethanol utilization is higher than in the other bottle experiments, likely caused by the addition of sulfate to the mineral stock solution. The aerobic treatment shows an earlier onset of succinate and lactate production than the anaerobic treatment. Addditionally, the specificity towards *n*-caproate is increased in the aerobic treatment compared to the anaerobic treatment. The pyrogluamate and pyruvate concentrations remained below detectable concentrations. The depicted values derive from daily or every other day conducted sampling, and the errors represent the standard error between biological triplicates.

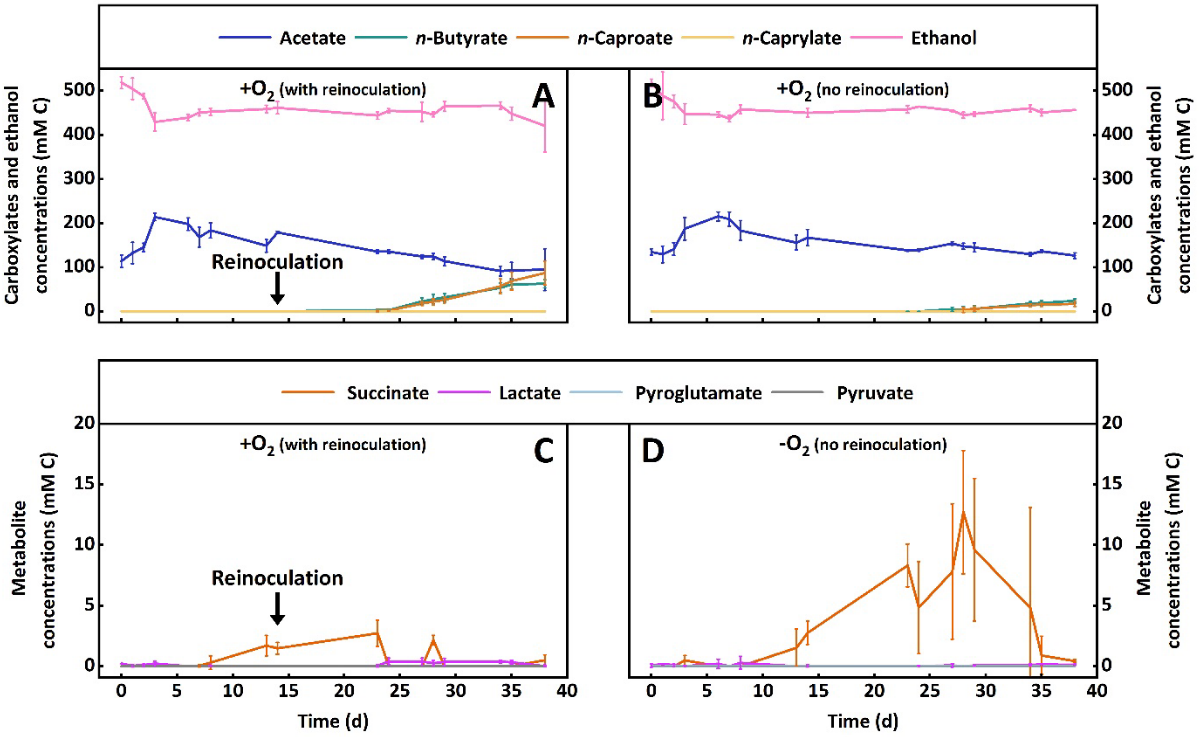


**Fig. S12.** Substrate, carboxylate (**A.** and **B.**), and intermediate metabolite (succinate, lactate, pyroglutamate, and pyruvate) (**C.** and **D.**) concentrations for the second additional bottle experiment throughout the 38-day experimental period. The treatment with anaerobic reinoculation is shown on the left figures (+$O_2$ with reinoculation) and the aerobic treatment without reinoculation on the right figures (+$O_2$ no reinoculation). The time of reinoculation is depicted with an arrow on day 14 (**A.** and **C.**). Both treatments show a more rapid ethanol utilization paired with acetate production during the first five days The succinate concentration remained lower, while the *n*-butyrate and *n*-caproate concentrations were progressing to higher values in the treatment with reinoculation, indicating that anaerobic species were promoted by the reinoculation and utilized succinate for chain elongation. The pyrogluamate and pyruvate concentrations remained below detectable concentrations. The depicted derive from from daily or every other day conducted sampling, and the errors represent the standard error between biological triplicates.

### Supplementary Tables

**Table S1. Bioreactor operation conditions, medium concentrations, and volumetric loading rate.** The wet working volume consists of the liquid-filled space used for microbial conversions. We included the filter modules in calculating the wet working volume as we noticed high biomass accumulation in this compartment, and their volume was non-negligible. The liquid level and, thus, the wet working volume in the bioreactor vessels rose when we attached the swan neck. Gas bubbles settled between the packing material in the AF reactor, displacing liquid, resulting in an uneven volume increase after the swan neck setup. On Day 963, we removed the filter modules from both bioreactor systems, reducing the total and wet working volume.

| Operating conditions | | | | | Substrate | | | Loading rate |
|---|---|---|---|---|---|---|---|---|
| Period | Time Start - End | Total volume | Wet working volume | HRT | Substrate concentration | Ethanol | Acetate | Total OLR |
| # | Day | L | L | Day | % | mM C | mM C | mmol C $L^{-1}$ $d^{-1}$ |
| **AF reactor** | | | | | | | | |
| **I** | 0 - 539 | 5.8 | 5.0 | 2.7 | 100 | 582.5 | 105.7 | 254.9 |
| **II** | 540 – 732 | 7.4 | 6.0 | 3.3 | 100 | 582.5 | 105.7 | 215.1 |
| **III** | 733 – 746 | 6.6 | 5.2 | 2.8 | 100 | 582.5 | 105.7 | 245.8 |
| | 747 – 766 | | | | 80 | 466.0 | 84.6 | 196.6 |
| | 767 – 861 | | | | 60 | 349.5 | 63.4 | 147.5 |
| **IV** | 862 – 971 | 6.6 | 5.2 | 2.8 | 60 | 349.5 | 63.4 | 147.5 |
| | 972 - 1019 | | | | 80 | 466.0 | 84.6 | 196.6 |
| **UASB reactor** | | | | | | | | |
| **I** | 0 - 539 | 5.8 | 5.8 | 3.3 | 100 | 582.5 | 105.7 | 215.1 |
| **II** | 540 – 732 | 7.4 | 7.4 | 4.2 | 100 | 582.5 | 105.7 | 167.9 |
| **III** | 733 – 746 | 6.6 | 6.6 | 3.7 | 100 | 582.5 | 105.7 | 186.0 |
| | 747 – 766 | | | | 80 | 466.0 | 84.6 | 148.8 |
| | 767 – 861 | | | | 60 | 349.5 | 63.4 | 111.6 |
| **IV** | 862 – 971 | 6.6 | 6.6 | 3.7 | 60 | 349.5 | 63.4 | 111.6 |
| | 972 - 1019 | | | | 80 | 466.0 | 84.6 | 148.8 |

**Table S2.** ***n*****-Caprylate productivities throughout the bioreactor operation.** The standard deviation reflects the variability of the volumetric *n*-caprylate production rate throughout the respective period.

| | | | AF reactor | UASB reactor |
|---|---|---|---|---|
| **Period** | **Time Start - End** | **Modification** | **Average volumetric *n*-caprylate production rate** | **Average volumetric *n*-caprylate production rate** |
| **#** | **Day** | | **mmol C $L^{-1}$ $d^{-1}$** | **mmol C $L^{-1}$ $d^{-1}$** |
| **I** | 0 - 539 | | 93 ± 43.1 | 73.5 ± 26.7 |
| **II** | 540 – 613 | Swan neck | 67.9 ± 31.8 | 90.1 ± 29.4 |
| | 614 – 644 | $N_2$ sparging headspace | 80.5 ± 35.5 | 43.4 ± 42.9 |
| | 645 – 700 | $N_2$ sparging medium tank | 43.7 ± 21.2 | 26.8 ± 23.2 |
| | 701 – 732 | Metal tubing | 47.4 ± 37.9 | 17.8 ± 21.7 |
| III | 733 - 861 | Filter module | 1.2 ± 2.1 | 2.5 ± 3.3 |

**Table S3. Influence of reactor modifications on oxygen concentrations.** The blue numbers represent the positions of the oxygen concentration measurements (Explanatory figure for **Table S3**). We measured the oxygen concentrations in the medium feed line (**1**), reactor inlet line (**2**), and at the start of the recycle line (**3**). The standard deviation reflects the variability of oxygen concentrations throughout the respective period.

| | | | AF reactor | | | UASB reactor | | |
|---|---|---|---|---|---|---|---|---|
| **Period** | **Time Start - End** | **Modification** | **1** | **2** | **3** | **1** | **2** | **3** |
| **#** | **Day** | | **mg L$^{-1}$** | | | **mg L$^{-1}$** | | |
| **End of I** | 515 - 539 | | 0.25 ± 0.05 | 0.44 ± 0.02 | 0.47 ± 0.05 | 0.28 ± 0.01 | 0.44 ± 0.02 | 0.48 ± 0.10 |
| **II** | 540 – 613 | Swan neck | 0.25 ± 0.01 | 0.32 ± 0.08 | 0.31 ± 0.01 | 0.13 ± 0.02 | 0.21 ± 0.01 | 0.42 ± 0.12 |
| | 614 – 644 | $N_2$ sparging headspace | 0.20 ± 0.07 | 0.20 ± 0.08 | 0.12 ± 0.01 | 0.12 ± 0.01 | 0.09 ± 0.01 | 0.13 ± 0.01 |
| | 645 – 700 | $N_2$ sparging medium tank | 0.04 ± 0.01 | 0.09 ± 0.01 | 0.10 ± 0.01 | 0.06 ± 0.01 | 0.09 ± 0.01 | 0.13 ± 0.01 |
| | 701 – 732 | Metal tubing | 0.03 ± 0.02 | 0.05 ± 0.02 | 0.05 ± 0.02 | 0.05 ± 0.01 | 0.05 ± 0.02 | 0.06 ± 0.03 |
| III | 733 - 861 | Filter module | 0.01 ± 0.01 | 0.02 ± 0.01 | 0.02 ± 0.01 | 0.02 ± 0.01 | 0.02 ± 0.01 | 0.03 ± 0.01 |

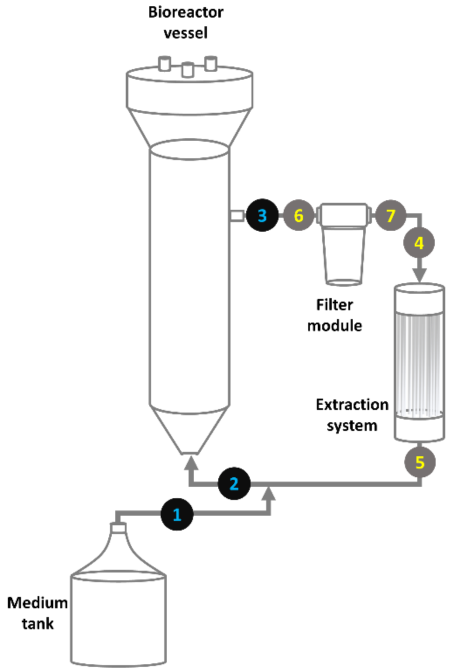


**Explanatory figure for Table S3.** The positions for the oxygen concentration measurements are highlighted. The blue numbers represent measuring points for long-term measurements. The aim of the long-term measurements was to elucidate the influence of reactor modifications on the oxygen concentrations. The yellow numbers represent short-term measurements (data given in the main text). We investigated the influence of the filter and extraction system on oxygen concentrations.

**Table S4. Unique proteins during high *n*-caprylate production.** We listed proteins uniquely found in the high *n*-caprylate production condition and listed the related microbial species, as well as the predicted catalyzed reaction. The proteins are ordered according to their taxonomic origin and from high to low abundance (intensity).

| Taxonomy | Protein name (Gene name) | Reaction | Intensity |
|---|---|---|---|
| **Actinomycetales** | | | |
| *Pseudoclavibacter caeni* | Alcohol dehydrogenase (adh2) | Propan-2-ol <=> Acetone | 8.2 |
| | Alcohol dehydrogenase (adhP) | Primary alcohol <=> Aldehyde | 7.5 |
| | Fumarate hydratase (fumC) | Malate <=> Fumarate + $H_2O$ | 7.4 |
| | 2-Methylcitrate dehydratase (prpD) | 2-Methylcitrate <=> But-2-ene-1,2,3-tricarboxylate + $H_2O$ | 6.9 |
| **Lactobacillales** | | | |
| Enterocaoccaceae | Acetate CoA-transferase (atoA) | Butanoyl-CoA + Acetoacetate <=> Butanoic acid + Acetoacetyl-CoA | 7.9 |
| **Bacteroidales** | | | |
| Bacteroidaceae | Fructose-bisphosphate aldolase (fdaB) | ß-D-Fructose 1,6-bisphosphate <=> Dihydroxyacetone phosphate + D-Glyceraldehyde 3-phosphate | 8.3 |
| | Isocitrate dehydrogenase [NADP] (icd) | Isocitrate <=> 2-Oxoglutarate + $CO_2$ | 8.1 |
| | Cysteine synthase (cysK) | *O*-Acetyl-L-serine + Hydrogen sulfide <=> L-Cysteine + Acetate | 6.8 |
| **Clostridiales** | | | |
| Clostridiales sp. | dUTP pyrophosphatase (dut) | dUTP + $H_2O$ <=> dUMP + Diphosphate | 8.4 |
| | Deoxyribose-phosphate aldolase (deoC) | 2-Deoxy-D-ribose 5-phosphate <=> D-Glyceraldehyde 3-phosphate + Acetaldehyde | 7.7 |
| | Acetyl-CoA carboxylase (accC) | ATP + Acetyl-CoA + $HCO_3$- <=> ADP + Orthophosphate + Malonyl-CoA | 7.6 |
| | Cysteine desulfurase (iscsS) | [Enzyme]-cysteine + L-Cysteine <=> [Enzyme]-S-sulfanylcysteine + L-Alanine | 7.0 |
| | Hydroxylamine reductase (hcp) | Ammonia + $H_2O$ <=> Hydroxylamine | 6.9 |
| | Nicotinamide-nucleotide ammidase (pncC) | Nicotinamide D-ribonucleotide + $H_2O$ <=> Nicotinate D-ribonucleotide + Ammonia | 6.8 |
| *Clostridium* sp. | Fructose-bisphosphate aldolase (fbaA) | ß-D-Fructose 1,6-bisphosphate <=> Dihydroxyacetone phosphate + D-Glyceraldehyde 3-phosphate | 7.6 |
| *Clostridium kluyveri* | Tryptophan synthase (trpB) | L-Serine + Indole <=> L-Tryptophan + $H_2O$ + D-Glyceraldehyde 3-phosphate | 7.9 |
| | valyl-tRNA synthetase (valS) | ATP + L-Valine + $tRNA^{Val}$ <=> AMP + Diphosphate + L-Valyl-$tRNA^{Val}$ | 7.8 |
| | 2-iminobutanoate/2-iminopropanoate deaminase (ridA) | 2-Iminobutanoate + $H_2O$ <=> 2-Oxobutanoate + Ammonia | 7.6 |
| | Pyruvate synthase (porA) | 2 Oxidized ferredoxin + 2-Oxobutanoate + CoA <=> 2 Reduced ferredoxin + Propanoyl-CoA + $CO_2$ + 2 $H^+$ | 7.4 |
| | Quinolinate synthase A (nadA) | Quinolinate + 2 $H_2O$ + Orthophosphate <=> Iminoaspartate + Dihydroxyacetone phosphate | 7.3 |

| | | | |
|---|---|---|---|
| | Serine hydroxymethyltransferase (glyA) | 5,10-Methylenetetrahydrofolate + Glycine + $H_2O$ <=> Tetrahydrofolate + L-Serine | 7.1 |
| | Phospho-2-dehydro-3-deoxy heptonate aldolase (aroA) | Phosphoenolpyruvate + D-Erythrose 4-phosphate + $H_2O$ <=> 2-Dehydro-3-deoxy-D-arabino-heptonate 7-phosphate + Orthophosphate | 7.1 |
| | UDP-glucose 4-epimerase (gale) | UDP-glucose <=> UDP-alpha-D-galactose | 6.7 |
| *Clostridium magnum* | Alanine--tRNA ligase (alaS) | ATP + L-Alanine + $tRNA^{Ala}$ <=> AMP + Diphosphate + L-Alanyl-tRNA | 6.8 |
| *Clostridium* sp. JN-9 | Acetyl-CoA synthetase (acsA) | ATP + Acetoacetate + CoA <=> AMP + Diphosphate + Acetoacetyl-CoA | 7.6 |
| *Oscillibacter ruminantium* | D-3-phosphoglycerate dehydrogenase (serA) | 3-Phospho-D-glycerate <=> 3-Phosphonooxypyruvate | 7.0 |
| *Oscillibacter valericigenes* | Phosphoserine aminotransferase (serC) | *O*-Phospho-L-serine + 2-Oxoglutarate <=> 3-Phosphonooxypyruvate + L-Glutamate | 7.8 |
| | Pyruvate-ferredoxin/flavodo xin oxidoreductase (nifJ) | Pyruvate:ferredoxin 2-oxidoreductase (CoA-acetylating), 2 Reduced ferredoxin + Acetyl-CoA + $CO_2$ + 2 $H^+$ <=> 2 Oxidized ferredoxin + Pyruvate + CoA Pyruvate:flavodoxin 2-oxidoreductase (CoA-acetylating), Pyruvate + CoA + Oxidized flavodoxin <=> Acetyl-CoA + $CO_2$ + Reduced flavodoxin | 7.7 |
| | Ribose 5-phosphate isomerase (rpiB) | D-Ribose 5-phosphate <=> D-Ribulose 5-phosphate | 7.7 |
| | Adenylosuccinate lyase (purB) | $N^6$-(1,2-Dicarboxyethyl)-AMP <=> Fumarate + AMP | 7.2 |
| *Sphaerochaeta pleomorpha* | Glutamate dehydrogenase [NADP] (gdhA) | L-Glutamate + $H_2O$ <=> 2-Oxoglutarate + Ammonia | 6.9 |
| **Methanobacteriales** | | | |
| *Methanobacterium congolense* | Thiamine thiazole synthase (thi4) | Glycine + [Protein]-L-cysteine <=> ADP-5-ethyl-4-methylthiazole-2-carboxylate + Nicotinamide + [Protein]-dehydroalanine + 3 $H_2O$ | 7.7 |
| | Orotidine 5-phosphate decarboxylase (pyrF) | Orotidine 5'-phosphate <=> UMP + $CO_2$ | 7.0 |
| *Methanobacterium paludis* | DNA-directed RNA polymerase (rboP) | ATP + $RNA_{(n)}$ <=> Diphosphate + $RNA_{(n+1)}$ | 7.1 |
| | Glutamyl-tRNA amidotransferase (gatA) | Glutaminyl-tRNA + L-Glutamate + Orthophosphate + ADP <=> L-Glutamyl-$tRNA^{Gln}$ + L-Glutamine + ATP + $H_2O$ | 7.0 |

### Supplementary Experimental procedures

**Bioreactor setup and operation.** We constructed and operated two upflow bioreactor systems. The setup and operation description apply to both bioreactor systems. The only constructional variation was that we filled packing material (Kaldnes K1 25 Liter, KoiCompetence, Witten, Germany) into the anaerobic filter (AF) reactor vessel while omitting packing material in the upflow anaerobic sludge blanket (UASB) reactor. An automated thermostat (CC 104A, Huber, Raleigh, NC, USA) recirculated heated distilled water through the heating jacket to maintain a fermentation broth temperature of 30 °C. We inserted a pH electrode (PL 80-325pH, SI Analytics, Weilheim, Germany) through a port on the top of the bioreactors. The pH electrode was attached to a controller (Alpha pH 800, Eutech Instruments, Singapore) with a connected peristaltic pump (Model 7542-12, Cole-Parmer, Vernon Hills, IL, USA). The controller automatically adjusted the fermentation broth to pH 5.5 by adding hydrochloric acid (3 M) through another port near the pH electrode. Another peristaltic pump (Model 07528-20, Cole Parmer) continuously supplied fresh medium kept at room temperature from a 5 L supply tank to each bioreactor at similar flow rates of 1.80 ± 0.04 L $d^{-1}$. The wet working volume of the AF reactor was smaller than that of the UASB reactor, resulting in a shorter hydraulic retention time of 2-7 – 3.3 d compared to 3.3 – 4.2 d (**Table S1**). A part of the fermentation broth exited the bioreactors through an overflow line into the effluent tank *via* a hose.

**Extraction system.** Products were extracted with a membrane-based liquid-liquid extraction system(*i.e.*, pertraction), as reported previously.[2, 3] The extraction system consisted of a forward and a backward membrane contactor (Liqui-Cel EXF-4 x 13 Membrane Contactor, 316L SS body, X50 membrane, 3M, Charlotte, NC, USA). Hollow-fiber membranes composed of polypropylene with a surface area of 8.1 m² each establish a liquid-liquid interface between two liquid streams inside the membrane contactors. While the forward membrane contactor creates a liquid-liquid interface between the fermentation broth and the hydrophobic solvent, the backward membrane contactor creates a liquid-liquid interface between the hydrophobic solvent and an alkaline extraction solution. In both cases, products transfer from one liquid stream to another *via* pertraction. A peristaltic pump (Model 7542-30, Cole-Parmer, Vernon Hills, IL, USA) recycled the fermentation broth between the bioreactor vessel and the forward membrane contactor through plastic tubing (Tygon® A-70-F, Masterflex, Gelsenkirchen, Germany) at a speed of ~7.0 L $h^{-1}$. Before entering the forward membrane contactor, the incoming fermentation broth was channeled through a 30 µm filter (GXWH20S, FXWSC filter, General Electric Company, Boston, MA, USA) with a liquid volume of 0.8 L to prevent fouling and associated plugging of the hollow fiber membranes.

The hydrophobic solvent was recycled (Model 7542-30, Cole Parmer) through the hollow fiber lumen sides (fiber interior) of the forward and backward contactor at a flow rate of 1.26 L $h^{-1}$ in the AF system and 1.10 L $h^{-1}$ in the UASB reactor. An alkaline extraction solution was recycled with a pump (Model 07528-30, Cole Parmer) between the shell side of the backward contactor and a 4 L stirred vessel at a flow rate of 5.1 L $h^{-1}$ in both bioreactor systems. The hydrophobic solvent comprised mineral oil with 30 g $L^{-1}$ tri-*n*-octylphosphine oxide (TOPO). Favorably higher-carbon-chain carboxylates extract through the liquid-liquid interface into the hydrophobic solvent. The buffered alkaline extraction solution consisted of distilled water with 30 g $L^{-1}$ sodium borate and 7 g $L^{-1}$ sodium hydroxide. During operation, a controller (Alpha pH 800, Eutech Instruments, Singapore) and a corresponding pump (Model 7542-12, Cole-Parmer, Vernon Hills, IL, USA) adjusted the pH of the alkaline extraction solution to pH 9 with the addition of sodium hydroxide (5 M). In the hydrophobic solvent, carboxylates appear in the protonated (acid) form. In the alkaline extraction solution, carboxylates are deprotonated (salt). The acid and salt form imbalance causes directed diffusion and accumulation of the carboxylates from the hydrophobic solvent into the alkaline extraction solution.

**Defined bioreactor medium.** We prepared the medium stock in a 10-L beaker by first adding 8-L tap water and then successively 300-mL mineral stock solution, 100-mL trace metal stock solution, 100-mL vitamin stock solution, 30 g sodium bicarbonate, 50 g 2-(*n*-morpholino)ethanesulfonic acid, 170 mL ethanol, and 30 mL acetate (except for periods with reduced the substrate concentration). Afterward, we filled the mixture to 10 L with tap water and shook the beaker in rotary movements to homogenize the ingredients. We adjusted the pH to 5.0 with sodium hydroxide or hydrochloric acid. The concentrations of mineral salts per Liter in the stock solution were:4 g sodium chloride; 5 g ammonium chloride; 5 g potassium chloride; 5 g potassium monophosphate; 10 g magnesium chloride; 2 g calcium chloride. The concentrations of trace metals per liter in the stock solution were: 2 g nitrilotriacetic acid; 1 g manganese sulfate; 0.8 g ammonium iron(II) sulfate; 0.2 g cobalt chloride; 0.2 g zinc sulfate; 0.02 g copper(II) chloride; 0.02 g nickel chloride; 0.02 g sodium molybdate; 0.02 g sodium selenate; 0.02 g sodium tungstate.

The concentrations of vitamins per liter in the stock solutiom were were: 0.01 g pyridoxine; 0.005 g thiamine; 0.005 g riboflavin; 0.005 g calcium pantothenate; 0.005 g thioctic acid; 0.005 g 4-amino benzoic acid; 0.005 g nicotinic acid; 0.005 g vitamin B12; 0.002 g d-biotin; 0.002 folic acid; 0.002 2-mercaptoethanesulfonic acid.

We did not add bromethanesulfonic acid or any other methanogen inhibitor. Also, we prepared the medium under ambient air without sterilization and kept the 10-L medium stock beaker in a 4 °C storage room until usage. When the 4-L medium supply tanks of the bioreactor systems were running empty, we filled them with fresh medium from the medium stock. The bioreactor medium supply tanks had no cooling system integrated, and we kept them at room temperature (~21 °C) during the operating period.

**Analytical procedures.** For the non-invasive dissolved oxygen concentration measurements, we used the OXY-4 SMA multi-channel oxygen meter (PreSens - Precision Sensing GmbH, Regensburg, Germany) with attached polymer optical fiber and oxygen sensor spots SP-PSt3-NAU-D5-YOP (PreSens - Precision Sensing GmbH). We measured the cell density ($OD_{600}$) with a NanoPhotometer NP80 (Implen, Westlake Village, CA, USA) at 600 nm with a path length of 0.67 mm. For the determination of both the ethanol and the simple carboxylate concentrations during bioreactor operation and batch experiments, we used an Agilent 7890B gas chromatograph (GC) (Agilent Technologies Inc., Santa Clara, CA, USA) equipped with a capillary column (DB Fatwax ultra inert 30 m × 0.25 mm × 0.25 µm, length × diameter × film thickness; Agilent Technologies Inc.) and an FID detector with hydrogen gas as mobile phase. The ramp temperature program and sample preparation protocol we used to quantify simple carboxylate concentrations corresponds to an earlier report with a slightly lower injection temperature of 200 °C, a detector temperature of 250 °C, and 100 times dilution of the extraction solution samples.[4] For ethanol quantification, we set a stable oven temperature of 40 °C for 4 min with a post-run temperature of 180 °C for 4 min, with the same injection and detector temperature as for the carboxylates. We determined the concentrations of succinate, lactate, pyroglutamate, and pyruvate with a Shimadzu LC20 high performance liquid chromatograph (HPLC) (Shimadzu, Kyoto, Japan) equipped with an Aminex HPX-87H column (Bio-Rad, Richmond, CA, USA), 5 mM sulfuric acid as mobile phase, and an RID-20A high-sensitivity refractive index detector (Shimadzu, Kyoto, Japan).[5] We set the acquisition program to a flow of 0.6 mL $min^{-1}$ with an oven temperature of 40 °C without ramp and 30 min per injection.

We defined the isotopic compositions of *n*-butyrate, *n*-caproate, and *n*-caprylate throughout the batch experiments with an Agilent 7890B/5977B gas chromatograph coupled with a mass spectrometer (GC/MS) (Agilent Technologies Inc., Santa Clara, CA, USA) which was equipped with a capillary column (DB Fatwax ultra inert 30 m × 0.25 mm × 0.25 µm, length × diameter × film thickness; Agilent Technologies Inc.) and an adjusted ramp temperature program (initial temperature of 40 °C for 3 min, a first ramp of 12 °C $min^{-1}$ from 40 °C to 160 °C, a second ramp of 20 °C $min^{-1}$ from 160 °C to 230 °C with 5 min hold) with hydrogen as carrier gas and electron ionization at 70 eV. After the batch experiments, we conducted the GC/MS measurements of produced carboxylates in one batch. Until then, we stored 500 µL of each sample at -20 °C. We used 50 µL of each stored sample for further preparation. The sample preparation involved transferring the liquid into Eppendorf PCR tubes, adjusting the pH below 2 with 10% phosphoric acid (v/v %), and adding 50 µL *n*-hexane as solvent. We shook the PCR tubes for 30 min, positioned horizontally, to increase the surface area between sampled liquid and solvent. Then, we placed the PCR tubes upright, waited for phase separation, and transferred the *n*-hexane-carboxylate mixture into glass micro-inserts (Rotilabo ND8 0.15 ml, Cal Roth, Karlsruhe, Germany) for small-volume GC/MS analysis.

**Stable isotope tracing.** We exported the total ion chromatograms (all *m/z* values included) of GC/MS measurements from the operating software of the system as .cdf files (OpenLab CDS version 2.7, Agilent Technologies Inc., Santa Clara, CA, USA). Next, we analyzed the extracted ion chromatograms (selected *m/z* ranges) with the Metabolomic Analysis and Visualization Engine (El-MAVEN v0.12.0) to quantify the $^{13}C$-isotopomer abundances of *n*-butyrate, *n*-caproate, and *n*-caprylate.[6-8] The observed fragmentation patterns allowed us to track the extent of labeling of multiple carbon-chain segments per analyzed carboxylate (**Fig. S1, S2**).

**Bottle experiments without stable-isotope tracing.** We conducted two additional bottle experiments in 200-mL serum bottles with a working volume of 25 mL and a headspace volume of 175 mL to complement the results of the stable-isotope tracing experiments by measuring succinate, lactate, pyroglutamate, and pyruvate as potential intermediate metabolites with the HPLC. For both experiments, we used the bioreactor medium with 3 g $L^{-1}$ bicarbonate, 5 g $L^{-1}$ 2-(*n*-morpholino)ethanesulfonic acid buffer, and added 0.5 g $L^{-1}$ L-cysteine hydrochloride. We derived the thawed inoculum from the filter module of the AF reactor on Day 701. When we removed the filter module, we filled the contained high-density biomass into 50-mL Falcon tubes, centrifuged them at 3450 × *g* (at 4 °C) for 15 min, discarded the supernatants, resuspended the emerged pellets in 10 mL sterile filtered tap water and stored the biomass in anaerobic serum bottles at -80 °C. We inoculated the bottles with 0.5 mL (2% (v/v) of the thawed biomass. Both bottle experiments were incubated at 30 °C without shaking and differed slightly in their experimental setup:

For the first additional bottle experiment, we exchanged magnesium chloride with 0.3 g $L^{-1}$ magnesium sulfate. We added 5 g $L^{-1}$ sodium chloride while the concentrations of ethanol and acetate were 240 mM C and 60 mM C, respectively. We incubated the bottles for 15 days. Following the stable-isotope tracing experiment, we set up one triplicate of bottles with aerobic and one triplicate of bottles with anaerobic starting conditions. For the second additional bottle experiment, we did not change the salt, ethanol (600 mM C), or acetate (100 mM C) concentrations. The experiment lasted 38 days; both triplicates had aerobic starting conditions. We reinoculated one of the triplicates on day 14 with 0.5 mL of additional inoculum.

***Clostridium kluyveri*** **isolate.** To isolate *C. kluyveri*, we used agar plates with a modified Reinforced Clostridial Medium. We prepared the medium according to previous reports, adding ethanol as selection pressure and adjusting the pH to 5.5 with 5 M hydrochloric acid.[9] After picking colonies, we incubated *C. kluyveri* at 30 °C in 50-mL serum bottles filled with 20-mL modified Reinforced Clostridial Medium. After a growth phase of one week, we extracted 1 mL of the cell culture for 16S rRNA gene sequencing. We physically lysed cells with a FastPrep-24™ 5G (MP Biomedicals, Irvine, CA, USA), following the instructions of the manufacturer (40 s at 6.0 m $s^{-1}$). Next, according to the corresponding protocol, we extracted DNA with a NucleoSpin® Microbial DNA Kit (Macherey-Nagel, Düren, Deutschland). We amplified the 16S rRNA gene with the universal primers 27F/1492R and sequenced the amplicon *via* Sanger sequencing. We trimmed the resulting genetic sequence, and the remaining 16S rRNA gene comprised 908 base pairs (63.4% completeness). After clarifying the taxonomy, we stored the cells in glycerol stock solutions at -80 °C until performing the batch experiments.

For the batch experiments, we thawed the glycerol stocks slowly on ice and precultured them two consecutive times in 100 mL serum bottles with a working volume of 20 mL. The headspace contained a nitrogen/carbon dioxide gas mixture (80/20% v/v), and the medium was modified Reinforced Clostridial Medium. We lowered the initial ethanol concentration to 7.89 g $L^{-1}$ (171.3 mM) for optimal growth for the batch experiments in pure culture. We adopted the acetate concentration according to the recipe, accounting for 2.16 g $L^{-1}$ (36.6 mM). We inoculated three replicate serum bottles with 1.2 mL of the second preculture to reach a calculated $OD_{600}$ of 0.003. The batch experiment lasted six days with an initial pH of 5.5. We sampled two to three times per day. From the samples, we measured concentrations of ethanol, acetate, *n*-butyrate, *n*-caproate, and *n*-caprylate.

**Microbiome analysis**

**Metagenomics.** To extract the DNA from the fermentation broth samples, we used a FastDNA™ SPIN Kit for Soil (MP Biomedicals, Irvine, CA, USA), following the protocol of the manufacturer except when stated. First, we pelleted the biomass with a centrifuge at 4 °C and 3500 × *g* for 15 min. Then, we decanted the supernatant and equally distributed 50 µg wet biomass into the Lysing Matrix E tube with phosphate buffer and MT buffer provided with the kit. Next, we physically lysed the cells with a FastPrep-24™ 5G bead beater (MP Biomedicals, Irvine, CA, USA), following the protocol instructions (40 s at 6.0 m $s^{-1}$). We repeated the lysing step one time to guarantee the rupture of difficult-to-lyse cells. Between lysing steps, we cooled the samples on ice for 5 min.

After extracting the DNA, we prepared the DNA library with the Ligation sequencing kit SQK-LSK109 for the first three samples and SQK-LSK114 (Oxford Nanopore Technologies Ltd., Oxford Science Park, UK) for the last sample.

Before loading the DNA into the sequencer, we measured the concentration of double-stranded DNA with a Qubit Flex Fluorometer (Invitrogen, Carlsbad, CA, USA) utilizing a dsDNA HS Assay-kit of the same manufacturer. We then sequenced the DNA with a MinION sequencer (Oxford Nanopore Technologies Ltd., Oxford Science Park, UK) with an R9.4.1 flow cell for the first three samples and an R10.4.1 flow cell for the last sample.

After sequencing, the basecalling was performed using guppy basecaller v6.4.6. The resulting fastq files were aligned against the NCBI-nr database (downloaded February 2023) using diamond blastx v2.1.6 (Buchfink et al, 2021) using the optional parameters '-c 1 -b 12 --outfmt 100 --long-reads -p 64'. The resulting daa files were meganized using the daa-meganizer included in MEGAN v6.24.20 using the parameters '-lcp 51 -lg' with the newest mapping file available from February 2022.[10] The meganized files were loaded into MEGAN in comparison mode, and the aligned bases were normalized for sample size. The microbial abundances (aligned bases) for all taxa in the samples were exported and visualized accordingly.

**Metaproteomics.** We collected samples from the AF and the UASB reactor during regular operation (a high volumetric *n*-caprylate production rate) and a forced production-rate crash (a low volumetric *n*-caprylate production rate). The AF reactor did not fully crash as intended, so we did not include the related metaproteomics results. Nevertheless, we processed the samples from both bioreactor systems together (including the mixed-standard sample). We renamed the samples with internal platform identifiers as follows:

- R02: AF reactor low production rate
- R03: AF reactor production rate
- R04: UASB reactor low production rate
- R05: UASB reactor production rate

We resuspended the precipitated protein pellets in a denaturation buffer (6°M urea, 2°M thiourea, 60°mM Tris pH 8.0). We mixed equal amounts of each sample to generate a standard sample, omitting the low production rate sample of the UASB reactor due limited amount. We digested 40°µg of the standard and 20°µg of each sample with Trypsin as previously described. After digestion, we purified the samples on stage tips using five C18 discs to ensure sufficient material for peptide binding.[11] As described elsewhere, we subjected the stage tips immediately to dimethylation labeling, with the standard sample labeled on two independent stage tips.[12] We checked for label incorporation with 3% (v/v) of the sample. Correct mixing was ensured by mixing 2.5% (v/v) of each sample and controlling the mixing of the tags by liquid chromatography-mass spectrometry (LC-MS). We prepared two mixes in a 1:1:1 (light:medium:heavy) ratio: Standard (L) / R03 (M) / R02 (H) and Standard (L) / R04 (M) / R05 (H). Next, we fractionated the peptide mixes using the Pierce high pH reversed-phase peptide fractionation kit (Thermo Fisher Scientific, Waltham, MA, USA). Each fraction was then purified on stage tips, as described above.

For the LC-MS/MS measurements, we eluted the peptides from the stage tips with 80 µL of acetonitrile (80% v/v) and formic acid (1% v/v). After acetonitrile evaporation, we concentrated the samples with a speed vac and injected 3 µg peptides into the LC-MS system. We measured the samples with an EASY-nLC 1200 (Thermo Fisher Scientific, Waltham, MA, USA) coupled with an Orbitrap Exploris mass spectrometer (Thermo Fisher Scientific, Waltham, MA, USA). The peptides were chromatographically separated using 75 µm (ID), 20 cm packed in-house with reversed-phase ReproSil-Pur 120 C18-AQ 1.9 µm resin (Dr. Maisch GmbH, Ammerbuch, BW, Germany). We eluted the peptides over 90 min using fraction specific gradient of solvent B (80% ACN in 0.1% formic acid) followed by a washout procedure. Finally, we acquired the MS1 spectra between 300-1,750°Thompson at a resolution of 120,000 with an AGC target set to custom (300% normalized AGC). The top 12 most intense ions were selected for HCD fragmentation with an NCE of 28 using a dynamic exclusion window of 60 s. The MS2 spectra were acquired at a resolution of 15,000 with the AGC target set to standard.

We processed the raw data obtained from the instrument with the MaxQuant software (version 1.6.17.0).[13] The protein sequence databases used for the database search consisted of the predicted protein sequences from the microbial genomic sequencing of each bioreactor (AF reactor = 154,614 proteins; UASB reactor = 131,587 proteins) and frequently observed contaminants (248 entries). We included the dimethyl labeling on peptide N-term and lysines to differentiate standard (Light = 28.0313001284 Da), L-cysteine-treated (Medium = 32.056407112 Da), and L-cysteine-untreated samples (Heavy = 36.0756702794 Da). A false discovery rate of 1% was required at the peptide and protein levels. A maximum of two missed cleavages was allowed, and full tryptic enzyme specificity was required. Carbamidomethylation of cysteines was defined as fixed modifications, while methionine oxidation and N-terminal acetylation were set as variable modifications. Match between runs was enabled. Quantification was performed using the raw protein intensity values and a minimum peptide count of two. All other parameters were left to MaxQuant default settings. The reported intensity values correspond to the MaxQuant raw protein intensity.

We mapped predicted protein sequences from both bioreactors using the eggNOG-mapper software (v.1.0.3) and associated database (v.4.5).[14] The resulting functional annotation was then formatted to use within the Perseus software (v. 1.6.14.0).[15] In Perseus, we filtered out reverse hits, contaminant hits, and proteins only identified *via* a modified peptide. Separate significance B tests were performed for each sample to identify the proteins that change significantly (FDR ≤ 0.1) in abundance between the high and low volumetric *n*-caprylate-production rate conditions. For reverse ß-oxidation, we manually picked the related proteins, as no KEGG pathway is available.

**Metabolomics.** After thawing the samples, we analyzed the intracellular and extracellular metabolite concentrations from extracts using reversed-phase ion-pairing liquid chromatography. The method included ultra-high-performance liquid chromatography (UHPLC) (Thermo Scientific DionexUltiMate 3000, Thermo Fisher Scientific, Waltham, MA, USA) with a high-resolution mass spectrometer (MS) (Thermo Scientific Q Exactive quadrupole-Orbitrap hybrid MS, Thermo Fisher Scientific, Waltham, MA, USA). We set the electrospray ionization to negative mode to ionize metabolites and separated them *via* an Acquity UPLC BEH C18 column (130 Å, 1.7 µm, 100 mm × 2.1 mm; pore size, particle size, diameter × length; Waters, Milford, MA) maintained at 25 $^{o}$C using the eluents as previously described.[16] The flow rate was 0.18 mL min$^{-1}$. The sample injection volume was 10 µL. We deduced the concentrations of intracellular metabolites by subtracting the extracellular concentrations from the combined intracellular and extracellular concentrations. We performed the UHPLC-MS data analysis using the Thermo Scientific Xcalibur software.

### Bioreactor Equations

**Wet working volume (L).** The addition of packing material in the AF reactor vessel caused unequal wet working volumes ($\boldsymbol{V_{wet,1-3}}$) between both bioreactor systems, while the total volumes remained identical. We noticed a high biomass accumulation in the filter module, which we added to the wet working volume calculation. The wet working volume of both bioreactor systems changed depending on the experimental period. The fermentation broth levels and wet working volumes rose after connecting the swan necks to the bioreactor vessels. Interestingly, the wet working volume disparity between the vessels with and without swan neck was unequal because gas bubbles were entrapped between the packing material, reducing the wet working volume further than only by liquid displacement.

Period I $$\boldsymbol{V_{wet,1} = V_{Vessel} + V_{Filter\ module}}$$ (**Eq. S1**)

Period II $$\boldsymbol{V_{wet,2} = V_{vessel\ with\ swan\ neck} + V_{Filter\ module}}$$ (**Eq. S2**)

Period III & IV $$\boldsymbol{V_{wet,3} = V_{vessel\ with\ swan\ neck}}$$ (**Eq. S3**)

With

**Anaerobic filter reactor**

$V_{Vessel}$ = 4.2 L $V_{Filter\ module}$ = 0.8 L $V_{vessel\ with\ swan\ neck}$ = 5.2 L

**Upflow anaerobic sludge blanket reactor**

$V_{Vessel}$ = 5 L $V_{Filter\ module}$ = 0.8 L $V_{vessel\ with\ swan\ neck}$ = 6.6 L

Where

$V_{Vessel}$ = wet volume of the bioreactor vessel, L

$V_{Filter\ module}$ = wet volume of the filter module, L

$V_{vessel\ with\ swansneck}$ = elevated wet volume of the bioreactor vessel with attached swan neck, L

**Total organic loading rate (mmol C L$^{-1}$ d$^{-1}$).** Substrate concentrations and the influent flow rate volume positively impact the total organic loading rate ($\boldsymbol{OLR}$), while increased wet working volume has a lowering influence. We normalized the substrate concentrations to mmol C by multiplication with the ethanol and acetate carbon numbers. The wet working volumes and substrate concentrations changed multiple times during the experimental periods, resulting in fluctuating organic loading rates (**Table S1**).

$$\boldsymbol{OLR = \frac{(C_{Ethanol} + C_{Acetate}) \times 2 \times f}{V_{wet,x}}} \qquad \textbf{(Eq. S4)}$$

Where

$C_{Ethanol}$ = concentration of ethanol in the influent, mM

$C_{Acetate}$ = concentration of acetate in the influent, mM

$f$ = influent flow rate, L d$^{-1}$

$V_{wet,x}$ = wet working volume of the system during period x, L

**Alkaline extraction solution volume on the day n (L).** To account for the dilution of accumulating carboxylates, we extrapolated the volume of the alkaline extraction solution on day n ($\boldsymbol{V_{ex,n}}$) throughout the experiment. The alkaline extraction solution had a predefined volume when set up. We calculated the base solution (sodium hydroxide, 5 M) supplemented and added it to the known volume of the alkaline extraction solution of the prior sampling date. We exchanged the alkaline extraction solution when the volume exceeded the capacity of its vessel. After exchanging the extraction solution, we set the volume to the predefined value. We did not include the volume of the extracted and accumulating carboxylates in the term, as tests showed a general agreement of the volume increase when only regarding the base solution addition.

$$\boldsymbol{V_{ex,n} = V_{ex,n-a} + \frac{W_{base,n} - W_{base,n-a}}{D_{base}}} \qquad \textbf{(Eq. S5)}$$

Where

$V_{ex,n}$ , $V_{ex,n-a}$ = volume of the alkaline extraction solution on the day n and n-a, L

$W_{base,n}$, $W_{base,n-a}$ = weight of the base supplementation bottle on the day n and n-a, g

$D_{base}$ = density of the base addition bottle, g L$^{-1}$

**Volumetric production rate (mmol C $L^{-1}$ $d^{-1}$).** The calculation of the volumetric production rate comprises a fermentation broth part (yellow) and an alkaline extraction solution part (blue). We assumed the part of the fermentation broth as previously reported.[17] We modified the previously noted part of the alkaline extraction solution to account for the change in volume and concomitant dilution of carboxylates over time. The previously reported alkaline extraction solution part included a fraction, with the nominator being the concentration of carboxylates on day n ($\boldsymbol{C_{ex,n}}$) subtracted by the concentration of carboxylates on day n-a ($\boldsymbol{C_{ex,n-a}}$). The calculation proceeded by multiplying the resulting concentration change by a fixed volume of the alkaline extraction solution ($\boldsymbol{V_{ex}}$). Here, we split and extrapolated $\boldsymbol{V_{ex}}$ into the volume of extraction solution on day n ($\boldsymbol{V_{ex,n}}$) and the prior sampling day ($\boldsymbol{V_{ex,n-a}}$) (**Eq. S5**). We multiplied both volumes with the respective concentrations ($\boldsymbol{C_{ex,n}}$, $\boldsymbol{C_{ex,n-a}}$) to deduce the changing carboxylate amount between day n and day n-a. Finally, we divided the produced carboxylates over time by the wet working volume during period x (**Table S1**) to yield the volumetric production rate.

$$\boldsymbol{Volumetric\ production\ rate} = \frac{\boldsymbol{1}}{\boldsymbol{V_{wet,x}}}\left[\frac{\boldsymbol{C_{fe,n}V_{wet,x}}}{\boldsymbol{HRT_n}} + \frac{\boldsymbol{C_{ex,n}V_{ex,n} - C_{ex,n-a}V_{ex,n-a}}}{\boldsymbol{t_n - t_{n-a}}}\right] \quad \textbf{(Eq. S6)}$$

Where

$V_{wet,x}$ = wet working volume of the bioreactor during period x, L

$C_{fe,n}$ = concentration of the carboxylate in the fermentation broth on the day n, mM C

$HRT$ = hydraulic retention time on the day n, d

$C_{ex,n}$, $C_{ex,n-a}$ = concentrations of the carboxylate in the extraction solution on days n and n-a, mM C

$V_{ex,n}$, $V_{ex,n-a}$ = volume of the stripping solution on days n and n-a, L

$t_n$, $t_{n-a}$ = the day of operation n and n-a, d

$a$ = sampling interval, d

**Substrate-into-*n*-caproate or *n*-caprylate conversion selectivities (%, specific product/total substrate).** We calculated the conversion efficiencies to *n*-caproate or *n*-caprylate based on the total carbon feed. The conversion efficiency is independent of the changing wet working volume, and therefore denotes the system performance with greater consistency throughout the experimental periods than the volumetric production rates.

$$\boldsymbol{S_{C6,C8}} = \frac{\boldsymbol{VVD_{C6,C8}}}{\boldsymbol{OLR}} \times \boldsymbol{100\%} \quad \textbf{(Eq. S7)}$$

Where

$VVD$ = volumetric production rate of *n*-caproate (C6) or *n*-caprylate (C8) on the day n, mmol C $L^{-1}$ $d^{-1}$

$OLR$ = total organic loading rate on the day n, mmol C $L^{-1}$ $d^{-1}$

**Mass spectrometry equations**

We performed the following equations for all isotopomers of analyzable carbon-chain segments of *n*-butyrate, *n*-caproate, and *n*-caprylate **(Fig. S1)**. For example, the six-carbon-chain segment of *n*-caproate or *n*-caprylate includes calculations regarding the light (M+0) isotope and all heavy isotopomers (M+1, M+2, .., M+6).

**Heavy isotope ratio (dimensionless).** We measured the abundance ratios between naturally occurring heavy isotopomers (M+1, M+2,.., M+x) and the more abundant, light isotopes (M+0) with unlabeled *n*-butyrate *n*-caproate and *n*-caprylate. The ratios measure the natural heavy isotope incorporation of *n*-butyrate, *n*-caproate, and *n*-caprylate. For example, if the two-carbon-chain segment of *n*-butyrate incorporates one $^{13}C$ isotope (M+1), the peak shifts from *m/z* 60 to *m/z* 61. The abundance ratio is the abundance of the *m/z* 61 peak divided by the more prominent *m/z* 60 peak. Crucially, it is impossible to conclude which carbon position is labeled by looking at only one of the segments of the carbon chain. The isotopomer M+1 means that one of the two carbon atoms in a chain is labeled with a $^{13}C$-carbon. To predict labeling positioning, multiple carbon-chain segments need to be compared.

$$\boldsymbol{R_{M+x,y} = \frac{nA_{M+x,y}}{nA_{M+0,y}}} \qquad \textbf{(Eq. S8)}$$

Where

$\boldsymbol{nA_{M+x,y}}$= natural abundance of the M+x isotopomer peak of carbon-chain segment $\boldsymbol{y}$ from the investigated unlabeled carboxylic acid, %

$\boldsymbol{nA_{M+0,y}}$ = natural abundance of the M+0 peak of carbon-chain segment $\boldsymbol{y}$ from the investigated unlabeled carboxylic acid, %

**Compensated relative labeling extent (%).** The uncompensated abundance of an M+x isotopomer during the labeling experiment includes the naturally occurring heavy isotope portion and the microbially facilitated labeling portion. As we aimed to investigate the true extent of microbially facilitated heavy isotope labeling of carboxylic acids during the batch experiments, we compensated for naturally occurring heavy isotopes. Therefore, we subtracted the naturally occurring heavy isotope portion using the formerly identified ratio ($\boldsymbol{lA_{M+0,y}} \times \boldsymbol{R_{M+x,y}}$), from the uncompensated abundance of the investigated isotopomer ($\boldsymbol{lA_{M+x,y}}$). We determined the subtracted part in relation to the M+0 peak of labeled carboxylic acids.

$$\boldsymbol{A_{M+x,y} = lA_{M+x,y} - (lA_{M+0,y} \times R_{M+x,y})} \qquad \textbf{(Eq. S9)}$$

Where

$\boldsymbol{lA_{M+x,y}}$= uncompensated abundance of the M+x peak of a carbon-chain segment $\boldsymbol{y}$ from the investigated labeled carboxylic acid during the batch experiment, %

$\boldsymbol{lA_{M+0,y}}$ = uncompensated abundance of the M+0 peak of carbon-chain segment $\boldsymbol{y}$ from the investigated labeled carboxylic acid during the batch experiment, %

$\boldsymbol{R_{M+x,y}}$ = natural heavy isotope ratio of isotopomer M+x in carbon chain segment $\boldsymbol{y}$, dimensionless

**Isotopomer ratio (dimensionless).** We divided the abundances of each isotopomer by the added abundances of all the isotopomers within that carbon-chain segment. The compensated isotopomer ratio normalizes the proportion of a specific isotopomer within its carbon-chain segment. For instance, the three-carbon-chain segment of each carboxylate studied has four isotopomer peaks (M+0, M+1, M+2, M+3). The isotopomer ratio shows in which ratio the M+1 isotopomer occurs in the three-carbon-chain segment. The addition of all isotopomer ratios of a carbon-chain segment gives 1.

$$\boldsymbol{IR_{M+x,y}} = \frac{\boldsymbol{A_{M+x,y,}}}{\sum_{x=0}^{y} \boldsymbol{A_{M+x,y}}} \qquad \text{(Eq. S10)}$$

Where

$\boldsymbol{A_{M+x,y,}}$= compensated abundance of the M+x peak within the carbon-chain segment, (dimensionless)

$\boldsymbol{y}$ = the carbon-chain segment number, (dimensionless)

**Total $^{13}$C, $^{12}$C incorporations (%).** Apart from the isotopomer distributions, we also calculated the total $^{12}$C / $^{13}$C incorporation (% mol/mol) into carboxylates, simplifying the understanding of the total fraction of $^{13}$C isotope incorporation. In the case of *n*-butyrate, we extrapolated the total incorporation by assuming the same difference of total labeling between the 3 and 4-carbon segments as between the two and three-carbon-chain segments (the full *n*-butyrate molecule was not analyzable due to overlap with other fragments (*n*-butyric acid in **Fig. S2A**).

$$\boldsymbol{I_{^{13}C}} = \sum_{x=0}^{y} \boldsymbol{IR_{M+x,y}} \times \frac{\boldsymbol{x}}{\boldsymbol{Z_{CA}}} \times \mathbf{100}\ \% \qquad \text{(Eq. S11)}$$

$$\boldsymbol{I_{^{12}C}} = \sum_{x=0}^{y} \boldsymbol{IR_{M+x,y}} \times \frac{\boldsymbol{Z_{CA}-x}}{\boldsymbol{Z_{CA}}} \times \mathbf{100}\ \% \qquad \text{(Eq. S12)}$$

Where

$\boldsymbol{IR_{M+x,y}}$ = isotopomer ratio of the M+x peak of the carbon-chain segment $\boldsymbol{y}$, dimensionless

$\boldsymbol{y}$ = the carbon-chain segment number. Here, we consider the highest measurable carbon-chain segments to investigate the total incorporation of isotopes. Therefore $\boldsymbol{y}$ is 6 (*n*-caproate) or 8 (*n*-caprylate); for *n*-butyrate, the four-carbon-chain segment was not analyzable due to overlap with other fragments, dimensionless

$\boldsymbol{x}$ **=** Number of incorporated $^{13}$C-carbon isotopes (M+0 = 0, M+1 = 1, ..)

$\boldsymbol{Z_{CA}}$ = total number of carbon atoms of the investigated carboxylic acid, dimensionless)
